# Isoleucine absence from adult human hemoglobin is mirrored in mosquito proteomes and limits malaria parasite growth

**DOI:** 10.64898/2026.09.23.753855

**Authors:** Leah Houri-Zeevi, Martin Kampmann, Manoj T. Duraisingh, Leslie B. Vosshall

## Abstract

Amino acid composition within a proteome is conventionally viewed as a property of individual protein function. However, specific nutrient scarcity has been shown to reshape which amino acids are encoded in the genome of microbes. Mosquitoes have fed on blood for 200 million years, relying on blood amino acids for reproduction. We wondered whether this ancient dependency has left an imprint on the mosquito proteome. Here we show a proteome-wide shift to lower isoleucine usage in blood-feeding mosquitoes, mirroring the known isoleucine deficiency of adult human hemoglobin. In comparing amino acid composition across the proteomes of blood-feeding and non-blood-feeding mosquitoes, we found that blood feeders show systematically lower isoleucine usage and that highly expressed gut enzymes that digest the blood meal are nearly isoleucine free. Analysis of 707 vertebrate genomes reveals that isoleucine deficiency in major hemoglobin subunits is shared among mammals. Specifically, all non-Malagasy primates, including humans, lack isoleucine in their adult hemoglobin subunits while retaining it in fetal and embryonic subunits. We find that two other major blood proteins, serum albumin and immunoglobulin G, also have reduced isoleucine levels (∼1.4%) compared to the human proteome-wide average of 4.38%. Finally, we reasoned that this widespread isoleucine restriction could serve to defend adult red blood cells against isoleucine-dependent blood parasites that feed on them. We tested this hypothesis with the malaria parasite *Plasmodium falciparum*, which is known to arrest growth in adult red blood cells when extracellular isoleucine is withdrawn. We found that neonatal red blood cells purified from umbilical cord blood, which contain isoleucine-rich fetal hemoglobin, permit growth without extracellular isoleucine. This work demonstrates that nutritional scarcity is reflected in the genomes of animals, and that amino acid depletion of host protein sequences may serve a protective function against blood parasites.

## INTRODUCTION

The genetic code is built from deoxyribonucleic acid (DNA), transcribed into chains of ribonucleic acid (RNA), then translated into sequences of amino acids to generate proteins, the main functional unit of the cell. Proteins across the tree of life are typically assembled from the same twenty amino acids, and the order and frequency with which a genome encodes each amino acid is among the most basic properties of its protein repertoire. This usage is generally understood as a compromise between the structural and functional demands that proteins must satisfy and the mutational forces that act on the underlying DNA. Implicit in this view is the assumption that amino acids are freely available, so that a genome can “draw” on any residue based solely on need. Amino acids, however, are not only the output of informational symbols from the genetic code but physical building blocks. They include essential nutrients that many organisms cannot synthesize and must instead extract from their diet. When an environment or a diet leaves one amino acid in chronic short supply, that scarcity can itself become an evolutionary force that reshapes the proteins a genome encodes.

In microbes, this imprint of diet was first recognized in individual highly abundant proteins when nutrients became scarce. Under sulfur limitation, the cyanobacterium *Calothrix* induces an alternative, sulfur-poor isoform of its most abundant light-harvesting protein, phycocyanin, replacing the sulfur-containing amino acids methionine and cysteine to spare the scarce nutrient^1^. Beyond individual proteins, the same pressure has been detected genome-wide in microbes across the world’s oceans. In a study of microbial gene sequences isolated from 746 seawater samples, paired with measurements of the water each sample came from, the strength of selection against amino acid changes in those genes correlated with nitrate concentration more closely than with temperature, depth, or any other variable measured. In low-nitrate water, the changes most strongly selected against were the ones that would have added nitrogen atoms to a protein. The effect was larger in highly expressed genes, likely because the cell must supply the extra nitrogen atoms in every copy of the protein it makes^2^.

Whether pressure imposed by the chronic scarcity of a single amino acid has ever reshaped the proteome of a multicellular animal has, to our knowledge, not been shown. Blood-feeding mosquitoes are an ideal model to study this question. Blood feeding is ancestral to the mosquito family and most extant species feed on mammalian blood^3^. The blood-feeding lifestyle is confined to females and is strictly required for reproduction, because the amino acids released from blood protein are used to synthesize the vitellogenin proteins that provision the female’s developing eggs^4^. Most mosquito larvae feed on microorganisms and decaying plant and animal material in their aquatic habitat^5^. Larval food is dilute and patchily distributed, so the adult female’s blood meal is the dominant concentrated source of protein available for egg production^6^. Yet blood is a nutritionally unbalanced meal. Its protein mass is dominated by hemoglobin, and neither the alpha nor the beta chains of adult human hemoglobin contain any isoleucine^7^, an absence that was previously noted as shared among several mammals^8^. A mosquito feeding on mammalian blood therefore consumes a meal rich in protein yet poor in isoleucine, an essential amino acid that insects cannot synthesize and must obtain from their diet. Consistent with this, isoleucine is a limiting factor for mosquito reproduction. Females fed on isoleucine-poor blood produce fewer eggs and supplementing isoleucine increases egg output^8,9^.

While females of most extant species of mosquitoes feed on mammalian blood for reproduction, three lineages in three genera – *Toxorhynchites*, *Topomyia*, and *Malaya –* have independently abandoned blood feeding altogether. A minority of other mosquito species feed exclusively on non-mammalian hosts whose hemoglobin retains normal isoleucine levels. In a companion paper, we generated chromosome-level reference genomes for species from all three non-blood-feeding mosquito lineages and from frog-blood-feeding *Uranotaenia lowii* mosquitoes^10^. The convergent release from mammalian blood protein dependency in the non-blood feeders allowed us to test whether the isoleucine-poor diet provided by mammalian blood has left any trace in the proteomes of blood-feeding mosquitoes.

Here we find that the mosquito’s long history of blood feeding is reflected in its proteome. Mosquitoes that feed on mammalian blood use less isoleucine in their proteome than their frog-blood-feeding and non-blood-feeding relatives. This difference is most pronounced in the highly abundant gut enzymes that digest the mammalian blood meal. This mirrors the observations in microbes that nutritional scarcity has the greatest impact on highly expressed proteins. We further find that isoleucine depletion is a shared feature of mammalian adult hemoglobin, not present in embryonic and fetal subunits and other globin genes. This depletion is also present in the two other major blood proteins, albumin and the constant chains of immunoglobulin G. We reasoned that this widespread isoleucine restriction could serve to defend adult red blood cells against isoleucine-dependent blood parasites that feed on them. We tested this hypothesis with the malaria parasite *Plasmodium falciparum*, which is known to arrest its growth in adult red blood cells when extracellular isoleucine is withdrawn^11^. We found that neonatal red blood cells purified from umbilical cord blood, which contain isoleucine-rich fetal hemoglobin, permit growth without extracellular isoleucine. This work demonstrates that nutritional scarcity is mirrored in the genomes of animals, and that amino acid depletion in protein sequence may serve as protection against parasites.

## RESULTS

### Mosquito blood feeding and human blood composition

Mosquitoes have fed on blood for approximately 200 million years (Fig. 1a) and host preference varies across mosquito genera and species. Most living mosquitoes feed on mammalian blood, while a minority specialize on other hosts including amphibians, reptiles, and birds^3^ (Fig. 1b). While females of some species can produce a first batch of eggs without a blood meal, only three mosquito lineages, *Toxorhynchites*, *Topomyia*, and *Malaya*, have abandoned blood feeding altogether (Fig. 1b).

**Figure 1.**
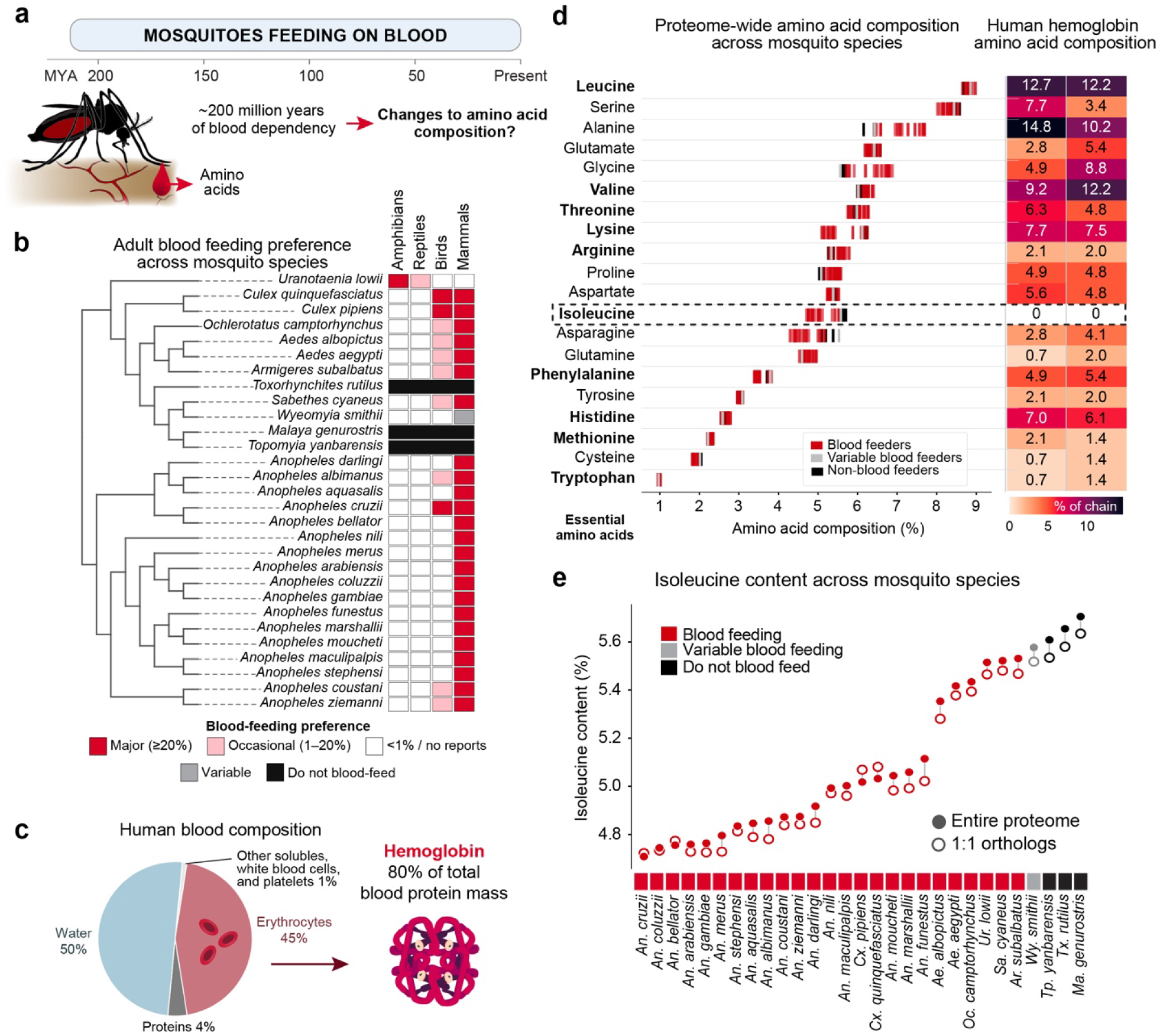
Blood-feeding mosquitoes encode less isoleucine across their proteomes. **a,** Timeline of mosquito blood feeding. **b,** Adult blood-feeding host preference across the mosquito species analyzed. Left, species tree. Right, reported preference for amphibian, reptile, bird, and mammal hosts: major host (≥20% of recorded meals, red), occasional host (1-20%, pink), variable blood-feeder *Wyeomyia smithii* (gray), non-blood feeder (black), and <1% or no reports (white). **c,** Composition of human blood by mass and contribution of hemoglobin to total blood protein mass. **d,** Left, proteome-wide amino acid composition across mosquito species. Rows are the twenty amino acids ordered by mean composition, and each tick is one species colored by feeding phenotype (blood feeding, red; variable blood feeding, gray; non-blood feeding, black). Essential amino acids are marked with an asterisk. Isoleucine (boxed) is the only amino acid whose usage is consistently lower in blood feeders compared to non-blood feeders. Right, amino acid composition of the human hemoglobin alpha and beta chains (% of chain); both contain zero isoleucine residues. **e,** Isoleucine content of each mosquito proteome, ordered from lowest to highest and colored by feeding phenotype as in **d**. Filled circles, entire proteome; open circles, 1-to-1 orthologs shared across all species.

Female mosquitoes depend on blood for their amino acid supply, and while mammalian blood is a protein-rich meal, it is very low in isoleucine, an essential amino acid for mosquitoes. By mass, human blood is roughly half plasma and half red blood cells. A single protein, hemoglobin, accounts for ∼98% of red blood cell protein^12^ and ∼80% of total blood protein mass (Fig. 1c)^13^. It has long been known that adult human hemoglobin is devoid of isoleucine^7^ and both the alpha and beta subunits entirely lack this essential amino acid (Fig. 1d). This means that female mosquitoes, which strictly depend on the amino acid nutrients in the mammalian blood meal, must reckon with a dietary isoleucine deficiency. We next asked if isoleucine scarcity is reflected in blood-feeding mosquito genomes.

### Reduced isoleucine usage in the proteomes of blood-feeding mosquitoes

We first tested proteome-wide amino acid usage in the protein-coding sequences of 29 mosquito genomes, spanning blood-feeding and non-blood-feeding species. While several amino acids varied in usage across the 29 genomes, isoleucine was the only amino acid whose usage was consistently reduced in blood feeders compared to non-blood feeders (Fig. 1d,e; Extended Data Fig. 1). Isoleucine usage also placed *Wyeomyia smithii*, a species with variable blood feeding, as an intermediate between blood-feeding and non-blood-feeding species (Fig. 1e). Isoleucine usage was reduced most in *Anopheles* and *Culex* mosquitoes and was higher in the two species that represent mosquito outgroups, the non-biting midge *Chironomus tepperi* and the biting midge *Culicoides brevitarsis* with isoleucine usage of 7.1% and 6.0%, respectively (Extended Data Fig. 1a).

We tested the difference in isoleucine levels across mosquitoes in a phylogenetic generalized least squares framework on the dated mosquito phylogeny, with the strength of phylogenetic signal (Pagel’s λ) estimated by maximum likelihood. Isoleucine usage was significantly lower in blood feeders than in the non-blood-feeding species and the variable blood feeder *Wyeomyia smithii* (β = −0.25, t₂₇ = −2.20, p = 0.037, λ = 0.88; phylogenetic simulation p = 0.033). The difference was present in the same direction when the three obligate non-feeders alone were compared with all other species (β = −0.23, t₂₇ = −1.96, p = 0.060; simulation p = 0.038).

Unlike human hemoglobin, amphibian hemoglobin retains normal isoleucine levels, so frog-blood-feeding mosquitoes do not face the isoleucine scarcity of mosquitoes that feed on mammalian blood. For example, in the African clawed frog *Xenopus laevis*, one of the major adult hemoglobin alpha subunits (UniProt: P02012) contains 5% isoleucine and a hemoglobin beta subunit (UniProt: P02132) contains 4.8% isoleucine. *Uranotaenia lowii*, the frog-blood-feeding mosquito, had higher isoleucine usage than most other blood feeders. Similarly, *Armigeres subalbatus* and *Sabethes cyaneus*, two mosquito species whose larvae consume a protein-rich diet by preying on other larvae in their aquatic habitat, also showed higher isoleucine usage than other blood feeders (Fig. 1e).

The reduction in isoleucine usage in blood-feeding mosquitoes was apparent whether we measured the full protein-coding repertoire, only proteins present in all species, or only 1-to-1 orthologs shared across all species (Fig. 1e, Extended Data^14^). This pattern cannot be explained simply by variation in genomic GC content (Extended Data Fig. 1b). Mutational and gene-conversion bias, measured as GC content at four-fold degenerate sites (GC4)^15,16^, correlated strongly with mosquito phylogeny and with the clades that contain the non-blood-feeding species, but did not separate blood feeders from non-blood feeders within a single clade (Extended Data Fig. 1c). The difference between blood feeders and non-blood feeders persisted when GC4 was included as a covariate, in both comparisons and in the same direction (β = -0.11, p = 0.011 for the three obligate non-blood feeders; β = -0.08, p = 0.075 including *Wyeomyia smithii*).

Isoleucine is an essential branched-chain amino acid, along with valine and leucine. While adult human hemoglobin lacks isoleucine, it contains ample valine (10.8%) and leucine (12.5%) residues. Despite the shared chemistry between valine, leucine, and isoleucine, isoleucine was the only branched-chain amino acid reduced in blood feeders. Valine and leucine content was elevated proteome-wide in most blood feeders compared to the non-feeders (Fig. 1d).

### Highly abundant blood-processing gut enzymes are isoleucine depleted

Selection to avoid a scarce amino acid is expected to fall most heavily on highly expressed proteins, which consume a disproportionate share of a cell’s amino acid budget, and to be strongest at the times when that resource is most limiting^1,17^. In the female mosquito this points to the proteins that are synthesized in bulk in the hours after a blood meal. These include gut proteases and peptidases that are expressed in the hours after a blood meal, and which digest blood protein into the free amino acids that are reassembled into yolk protein. Because this machinery is transient and is never packaged into the eggs, a female that built it out of isoleucine would be spending a scarce, egg-limiting resource on tools rather than on offspring.

We examined all three tissues that female *Aedes aegypti* use to process or benefit from a blood meal, fat body, ovary, and midgut, to ask if there is a selective reduction in transcriptionally-weighted isoleucine usage after blood feeding (Fig. 2a,b; Extended Data Figs. 2-4). For each, we used published datasets of the transcriptional response in the tissue after blood feeding^10^. We weighted amino acid usage by the relative share and length of each transcript, using transcription as a proxy for protein production. We found that only the midgut showed selective and significant isoleucine usage reduction.

**Figure 2.**
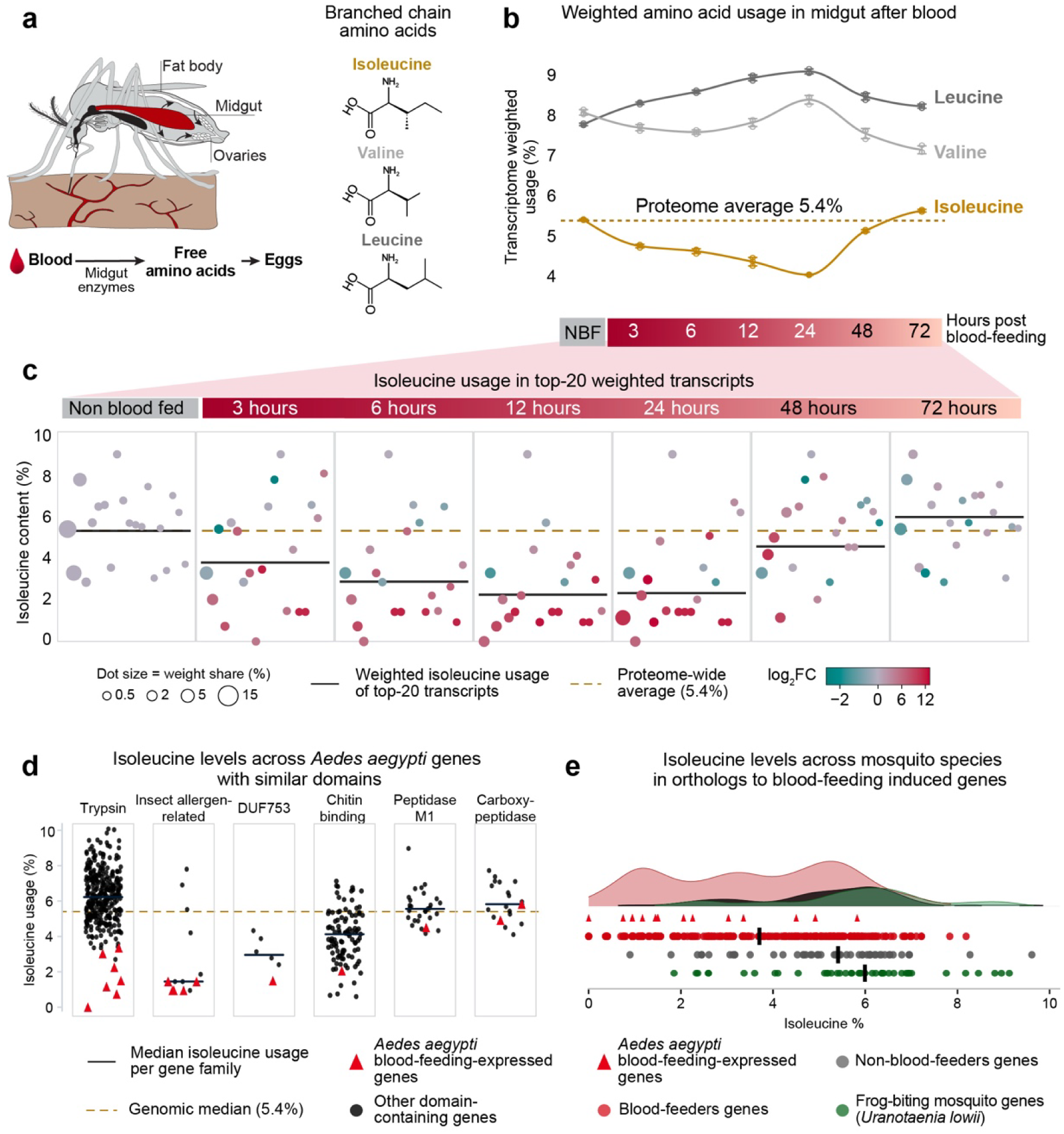
Isoleucine depletion is most pronounced in the blood-activated digestive enzymes of the female midgut. **a,** Left, tissues of the female *Aedes aegypti* involved in processing and using the blood meal. Right, structures of the three branched-chain amino acids. **b,** Transcriptionally-weighted usage of isoleucine (gold), valine and leucine (gray) in the female midgut of non-blood-fed females (NBF) and at 3, 6, 12, 24, 48 and 72 h after a blood meal. Weighted usage combines the amino acid composition of each protein with the relative share and length of its transcript. Dashed line, proteome-wide average isoleucine usage (5.4%). Points denote the mean. **c,** Isoleucine content of the twenty transcripts contributing most to weighted amino acid usage at each time point. Each circle is one transcript, sized by its share of the total transcriptional weight and colored by log2 fold change relative to the non-blood-fed midgut. Black line, weighted isoleucine usage of the top 20 transcripts; gold dashed line, proteome-wide average (5.4%). **d,** Isoleucine usage of blood-feeding-expressed midgut genes (red triangles) compared with all other *Aedes aegypti* genes carrying the same protein domain (black circles), shown separately for each domain family. Black line, median isoleucine usage per family; gold dashed line, genomic median (5.4%). **e,** Isoleucine content of the orthologs of the blood-feeding-expressed *Aedes aegypti* midgut genes across mosquito species. Density curves and individual genes are shown for blood feeders (red), non-blood feeders (gray) and the frog-blood-feeding *Uranotaenia lowii* (green); vertical bars denote group medians.

In the fat body, which uses amino acids that are digested in the midgut, the transcriptional response to blood was driven largely by vitellogenin expression (Extended Data Fig. 3). Vitellogenins form the egg yolk in the ovary and are massively upregulated after a blood meal and become the most abundant proteins in the fat body. In the main mosquito vitellogenin genes, Vitellogenin A1, B, and C, isoleucine is underrepresented, but so are the other two branched-chain amino acids, valine and leucine. In Vitellogenin A1, isoleucine accounts for 3.2% of residues compared to a proteome average of 5.4%, valine accounts for 4.9% compared to the 6.1% proteome average, and leucine accounts for 5.2% compared to a proteome average 8.7%. The reduced weighted isoleucine usage in the fat body therefore reflects generally lower branched-chain amino acid usage rather than a specific isoleucine reduction. Similarly, the ovary did not show a selective isoleucine reduction. We found that at 48 hours after feeding, there was a small reduction in both isoleucine and leucine usage driven largely by vitelline membrane proteins (Extended Data Fig. 4).

In contrast, the midgut showed a strong, significant isoleucine-specific reduction that was mirrored by elevated usage of valine and leucine (Fig. 2b). This reduction in isoleucine usage was most pronounced at 3, 6, 12, and 24 hours after feeding and returned to baseline by 48 hours (Fig. 2b), a time course that coincides with the heavy translational burden of the gut following a blood meal^18^. Examining the top 20 genes contributing to the weighted amino acid usage per timepoint revealed multiple blood-activated gut enzymes that are strongly, and in some cases completely, depleted of isoleucine. This included the 245-amino acid blood-feeding-specific Chymotrypsin-2 (*AAEL001690*) protease with zero isoleucine residues (Fig. 2c, Extended Data Fig. 2). Isoleucine depletion in these blood-activated midgut genes was pronounced even relative to other *Aedes aegypti* genes that share the same domains as the enzymes that are expressed in the gut after feeding on blood such as trypsins, peptidases, and carboxypeptidases (Fig. 2d).

Finally, among the blood-activated midgut genes shared across mosquito species, both non-blood-feeding mosquitoes and the frog-blood-feeding *Uranotaenia lowii* encoded more isoleucine residues in the direct orthologs of these isoleucine-depleted genes (Fig. 2e). A subset of the most highly expressed blood-activated midgut transcripts is specific to blood-feeding mosquitoes^10^ (Fig. 2e).

### Isoleucine depletion in adult mammalian hemoglobin

The absence of isoleucine from adult human hemoglobin was first reported nearly 70 years ago from amino acid composition analysis of purified protein before the protein’s amino acid sequence had been determined^7^. We note that the absence of isoleucine from adult human hemoglobin is a statistical anomaly. The mean proteome-wide human isoleucine frequency is 4.38%. Adult hemoglobin is a tetramer of two 141-residue alpha subunits and two 146-residue beta subunits. Chains of that combined length would be expected to contain no isoleucine with a random probability of 2.6 x 10^-6^ (0.9562^141+1^^46^). Examination of other oxygen-carrying globins, such as human myoglobin, reveals normal isoleucine levels. Human myoglobin contains 8 isoleucine residues in 154 amino acids, or 5.2% isoleucine composition.

The absence of isoleucine from adult hemoglobin was previously noted in several mammalian species^8^ but there has not been a comprehensive analysis of isoleucine usage in hemoglobin across the vertebrate tree of life. We therefore quantified isoleucine usage in hemoglobin subunits across 707 vertebrate genomes. We found that the percent of isoleucine encoded in the hemoglobin proteins dropped sharply in the two major adult mammalian subunits, alpha and beta. In contrast, isoleucine usage was generally higher in the non-major alpha or beta mammalian subunits and in non-mammalian hemoglobins, which used isoleucine at ∼3-5% across fish, amphibians, reptiles, and birds (Fig. 3a-c; Extended Data Fig. 5). In the major beta subunit, the loss of isoleucine was mirrored by an increase of valine, the structurally conservative replacement (Fig. 3c). Some mammalian lineages, including rodents and marsupials, carry isoleucine-containing hemoglobin. Other lineages, including most primates and multiple bat species, contain no isoleucine residues in any adult subunit.

**Figure 3.**
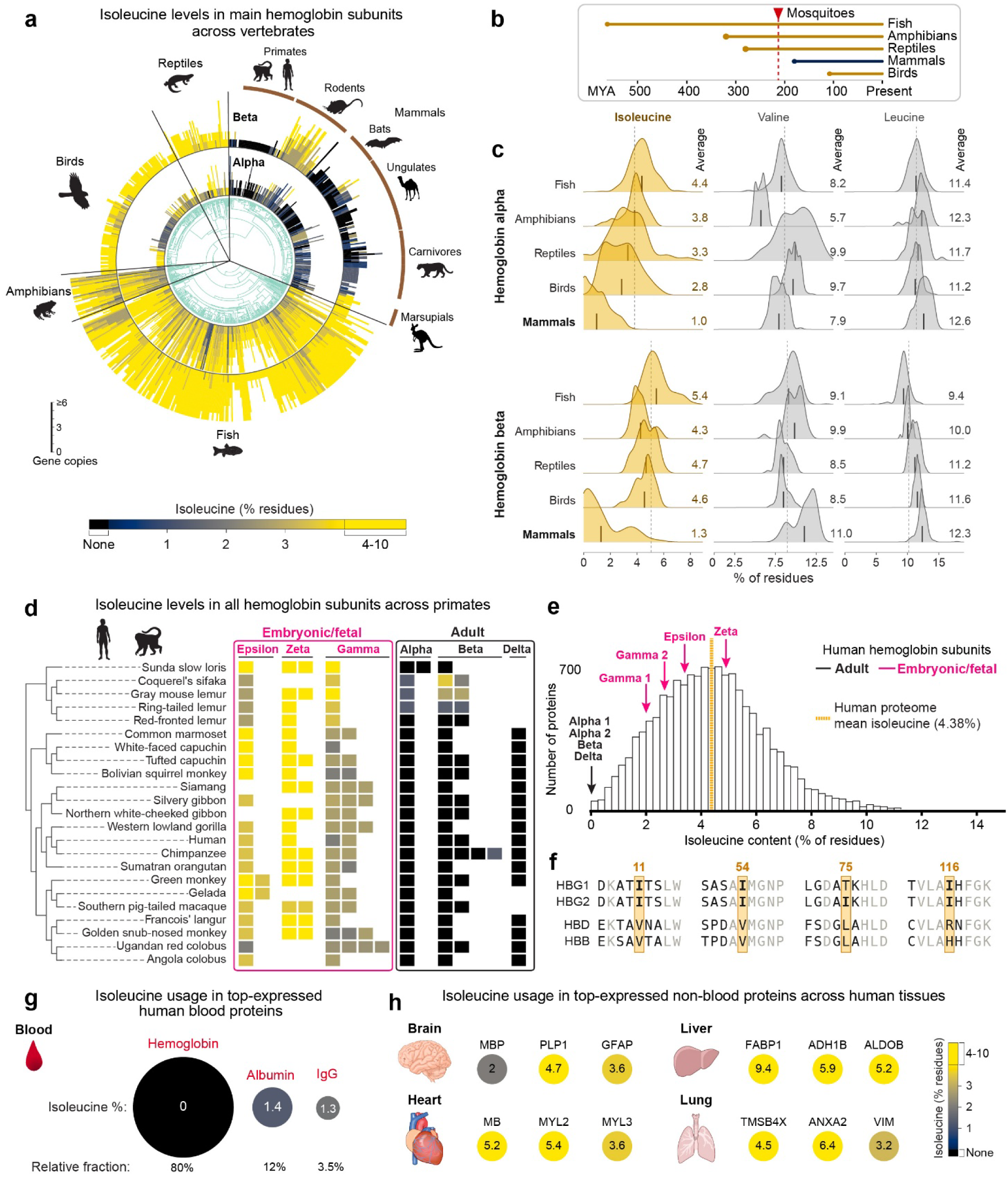
Isoleucine depletion is an adult-specific and conserved feature of mammalian blood proteins. **a,** Isoleucine content of the main hemoglobin subunits, alpha (inner ring) and beta (outer ring), across 658 species drawn from analysis of 707 vertebrate genomes, mapped onto the vertebrate phylogeny. The remaining 49 species are absent from the species-level timetree. Bars are colored by isoleucine content (% of residues), with black denoting no isoleucine residues. Major vertebrate and mammalian groups are indicated. **b,** Timeline of the origin of each vertebrate class, with the emergence of mosquitoes indicated (red, ∼200 million years ago). **c,** Distribution of isoleucine (gold), valine and leucine (gray) content in the hemoglobin alpha (top) and beta (bottom) subunits by vertebrate class. Vertical bars denote the class mean, given at right; dashed line, mean across all vertebrates. **d,** Isoleucine content of all hemoglobin subunits across primates, grouped by the developmental stage at which each subunit is expressed: embryonic and fetal (epsilon, zeta, gamma; magenta boxes), and adult (alpha, beta, delta; black box). Colored as in **a**. **e,** Distribution of isoleucine content across human proteins. Arrows mark the hemoglobin subunits: the adult alpha, beta and delta subunits at 0% (black), and the embryonic and fetal subunits (magenta). Orange dotted line, human isoleucine proteome average (4.38%). **f,** Alignment of the human fetal (HBG1, HBG2) and adult (HBD, HBB) hemoglobin subunits at the four positions occupied by isoleucine in the gamma subunits. **g,** Isoleucine usage in the three most abundant human blood proteins. Circle size denotes the share of total blood protein mass (percentage is given below each circle) and circle color and the value inside denote isoleucine content (% of residues), colored as in **h**. **h,** Isoleucine usage in the most highly expressed non-blood proteins of brain, heart, liver and lung, colored by isoleucine content (% of residues).

We next focused on primates, in which the developmental expression of hemoglobin subunits is best understood^19^. All three adult subunits, alpha, beta, and delta, contain zero isoleucine residues, whereas the fetal and embryonic subunits use isoleucine at normal levels (Fig. 3d). The primates of Madagascar are an exception to the primate rule of zero isoleucine residues in adult hemoglobin as several species encode a small number of isoleucine residues in their adult alpha and beta subunits (Fig. 3d). Against the full distribution of isoleucine usage among human proteins at least 120 amino acids long (Fig. 3e), we found that the adult hemoglobin subunits are clear outliers at 0% isoleucine, while the embryonic and fetal subunits fall within the typical proteome-wide range (Fig. 3e). Outside of hemoglobin, only 44 other proteins of the 14,296 in the human proteome (0.3%) in the human proteome lack isoleucine completely (Extended Data^14^). At the four isoleucine positions in the main fetal gamma hemoglobin subunit (HBG2), two of the corresponding adult sites are replaced by valine and one by leucine; both of which are the conservative replacements (Fig. 3f). The fourth isoleucine is replaced by arginine or histidine, which are non-conservative replacements.

### Other major human blood proteins are isoleucine deficient

We next asked whether isoleucine depletion is a hemoglobin-specific peculiarity or whether it is a shared property of other proteins with high expression in blood specifically or proteins with high expression in any tissue. To do this we searched the protein abundance database PaxDb for isoleucine content of highly expressed proteins across human tissues. The outcome of this analysis was that isoleucine deficiency is a feature of highly expressed blood proteins. The second and third most abundant blood proteins after hemoglobin are albumin, which transports fatty acids, hormones, and drugs in the blood plasma and helps maintain blood osmotic pressure^20^, and immunoglobulin G (IgG), the most abundant antibody in blood^21^. Albumin accounts for ∼12% of blood protein mass and IgG for ∼3.5%^13^, and together with hemoglobin they are estimated to make up about 95% of the total protein content in blood. We find that albumin and IgG also show reduced isoleucine levels (Fig. 3g). Human albumin encodes 8 isoleucine residues in a 585 amino acid mature protein (1.4% isoleucine). An assembled IgG encodes 10-12 isoleucine residues across its constant secreted chains totaling 866-874 residues (1.2-1.4% isoleucine). By contrast, the most highly abundant non-blood proteins in brain, heart, liver, and lung showed normal isoleucine usage overall. We conclude that isoleucine depletion is specific to blood rather than a general bias in translation or amino acid usage (Fig. 3h).

### *Plasmodium falciparum* escapes growth arrest in isoleucine-containing neonatal red blood cells

Finally, we hypothesized that the depletion of isoleucine from adult mammalian hemoglobin may represent an evolutionary defense against blood-borne pathogens. *Plasmodium falciparum*, the malaria parasite, relies heavily on hemoglobin catabolism as a primary amino acid source and digests hemoglobin in a dedicated food vacuole after entering the red blood cell^11,22^. We find that isoleucine is the third most used amino acid in the *Plasmodium falciparum* proteome (Fig. 4a). Yet, adult human hemoglobin contains no isoleucine and the parasite must acquire it from host plasma, where it circulates as a free solute at approximately 32-90 µM^23^. To reach that pool the parasite establishes new permeability pathways (NPPs) in the red blood cell membrane, which allow uptake of small solutes from outside the cell^23^. Consistent with this dependency, removal of isoleucine from culture medium has been shown to stall *Plasmodium falciparum* growth and induce drastically slowed development^11,24^.

**Figure 4.**
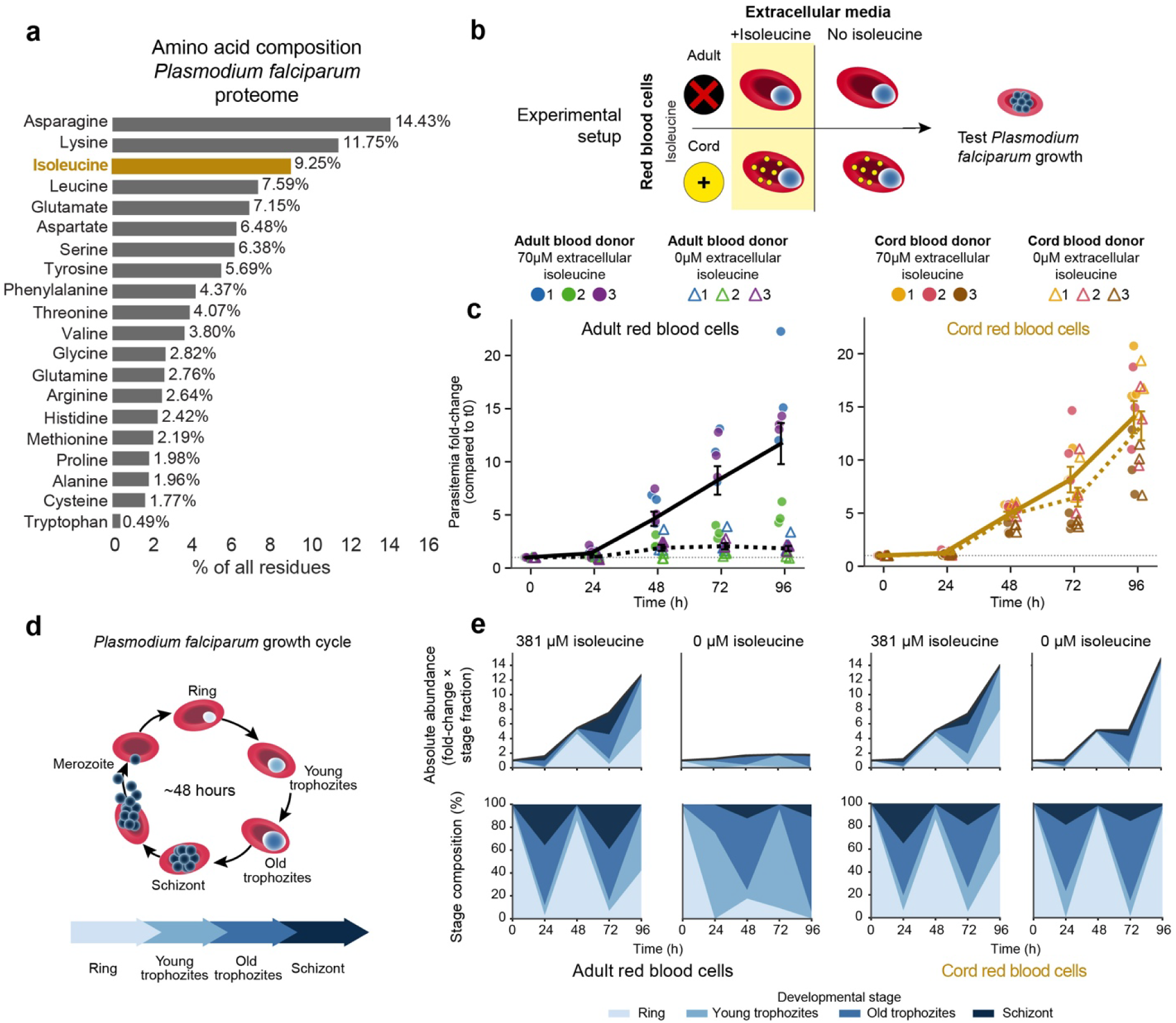
Isoleucine withdrawal arrests *Plasmodium falciparum* in adult but not fetal-hemoglobin-containing red blood cells. **a,** Amino acid composition of the *Plasmodium falciparum* proteome, ranked by relative residue levels (% of all residues). Isoleucine (gold) is the third most used amino acid in the parasite proteome. **b,** Experimental setup. *Plasmodium falciparum* was cultured in adult or cord (fetal hemoglobin-containing) red blood cells, in media with or without isoleucine, and parasite growth was measured. **c,** Parasitemia fold change relative to t0 in adult (left, black) and cord (right, gold) red blood cells over 96 h, in the absence of extracellular isoleucine (open triangles, dotted line; 0 µM) or at physiological levels (filled circles, solid line; 70 µM). Individual points denote *n* = 3 independent blood donors per blood type, colored by donor; lines denote the mean and error bars s.e.m. Dotted horizontal line indicates no change relative to t0 (fold change = 1). **d,** The ∼48-hour intraerythrocytic growth cycle of *Plasmodium falciparum*, progressing through ring, young trophozoite, old trophozoite, schizont and merozoite stages. **e,** Parasite stage distribution over 96 h in adult and cord red blood cells, at 381 µM and 0 µM extracellular isoleucine. Top, absolute abundance of each stage (parasitemia fold change x stage fraction); bottom, stage composition (% of parasites).

The embryonic and fetal subunits of hemoglobin, which retain isoleucine, are largely shielded from *Plasmodium*. Following the genetic switch to adult hemoglobin subunits that takes place around birth, fetal hemoglobin is only present in blood for the first few months of the newborn’s life^19,25^. During *in utero* development, the placenta limits parasite access to the bloodstream of the developing fetus^26,27^.

*Plasmodium falciparum* can be maintained in continuous *in vitro* culture in human erythrocytes using RPMI 1640 medium, a system established fifty years ago by William Trager and James B. Jensen at Rockefeller University in the same building where the present study was conducted^28^ and it remains standard practice today. RPMI 1640 contains 381 µM isoleucine, far above physiological plasma levels. Using such high-isoleucine culture systems, earlier studies suggested that fetal hemoglobin in fact provides a protective effect against *Plasmodium* parasites^29^. This view has since been overturned by works showing that *Plasmodium* parasites grow equally well in adult and fetal hemoglobin-containing red blood cells under conditions of high isoleucine^30,31^.

Together, we propose that the depletion of isoleucine specifically from adult hemoglobin restricts intracellular blood parasites. This predicts that when extracellular isoleucine is scarce, fetal hemoglobin, by virtue of its own isoleucine content (4 of 146 residues in the major gamma subunit, 2.7%) and despite not being the parasite’s usual protein source, should allow normal parasite growth rather than restrict it.

To determine whether isoleucine derived from hemoglobin digestion alone could sustain parasite growth in the absence of exogenous isoleucine, we cultured *Plasmodium falciparum* in adult red blood cells or in cord blood red blood cells across extracellular isoleucine concentrations of 0, 10, 20, 70 and 381 µM in three independent donors each (Fig. 4b). Cord blood here refers to human neonatal blood collected from the umbilical cord at delivery, in which fetal hemoglobin accounts for roughly 70 to 80% of total hemoglobin at term^25,30,32^. Accordingly, we analyzed the presence of fetal hemoglobin in red blood cells obtained from three adult donors and cord blood samples from three donors and found that fetal hemoglobin was absent or nearly absent in all adult red blood cells and present in nearly all cord red blood cells (Extended Data Fig. 6).

Consistent with prior reports^11,33,34^, parasites cultured in adult blood exhibited near-complete growth arrest upon isoleucine withdrawal, growing 1.7-fold over 96 hours without isoleucine compared with 10.3-fold in the presence of 70 µM isoleucine (Fig. 4c, Extended Data Fig. 7). In stark contrast, parasites cultured in cord blood grew comparably with 70 µM and 0 µM extracellular isoleucine (13.3-fold and 12.6-fold, respectively). Isoleucine withdrawal reduced growth ∼6-fold in adult blood (paired *t*-test on log₂ fold change, *t_2_* = 11.4, p = 0.008, *n* = 3 donors) but had no detectable effect in cord blood (*t_2_* = 1.9, p = 0.20), and a linear mixed-effects model over the full time-course confirmed that this dependence of the isoleucine effect on red-blood-cell category was significant (blood type × isoleucine × time interaction, p < 0.001). Parasites grown in 10, 20 or 381 µM isoleucine media followed the same growth trajectory as the 70 µM isoleucine condition (Extended Data Fig. 7, Extended Data^14^).

*Plasmodium falciparum* completes a growth cycle inside the red blood cell in about 48 hours, passing through a well-characterized series of stages known as the intraerythrocytic developmental cycle (Fig. 4d). In adult blood without isoleucine, progression through this cycle was severely impaired, with only a small proportion of parasites advancing to late stages and at a markedly reduced rate (Fig. 4e, Extended Data Fig. 8). Parasites in cord blood progressed comparably at 381 µM extracellular isoleucine concentration and under complete isoleucine withdrawal.

## DISCUSSION

Our findings suggest that over ∼200 million years of blood feeding, mosquitoes have become both witnesses to and participants in an evolutionary contest over blood, and that their proteomes carry a record of this conflict. This provides, to our knowledge, the first demonstration that dietary pressure is reflected in proteome-wide amino acid usage in a multicellular animal.

The contest over blood is fundamentally tripartite. The female mosquito needs blood to reproduce; the malaria parasite needs the mosquito and vertebrate blood to complete its life cycle; and the vertebrate host, which likely needs none of this, is nonetheless the reservoir from which infected blood is drawn back into mosquitoes. *Plasmodium* parasites can pass between hosts only through blood and must therefore remain in blood. Isoleucine scarcity is a thread that runs through all three players: a limiting nutrient for the mosquito, an essential and hard-won resource for the parasite, and as we hypothesize, a protective feature of the host tissue that parasite and mosquito compete over.

A plausible selective pressure for the mosquito shift in isoleucine usage is already established. Isoleucine is a limiting resource for females feeding on isoleucine-poor mammalian blood, and supplementing a blood meal with isoleucine increases the number of progeny a female produces^8,9^. The depletion we observe is most dramatic in the blood-activated digestive enzymes of the midgut and peaks during the window of heaviest post-blood-meal translation, when demand for amino acids is greatest. Building highly induced proteins from a scarce amino acid could stall rapid, high-demand translation^35^. The reduced isoleucine in these strongly induced proteins may therefore originate in selection on translational efficiency as much as on the total isoleucine budget.

Does this shift reflect selection on isoleucine itself, or is it a neutral consequence of genome-wide base composition? Isoleucine is encoded by AT-rich codons, so a rise in GC content would lower its usage passively. Several observations argue against a purely neutral explanation. Genomic GC content does not track the blood-feeding phenotype, mutational bias does not separate blood feeders from non-blood feeders within a clade, and the depletion is most pronounced within a single genome in the blood-activated gut enzymes where base composition is constant. A neutral contribution to the proteome-wide signal cannot be excluded. An intriguing possibility remains that mutational biases themselves might be tuned by selection to economize on a limiting resource or drive a shift in diet to accommodate changing proteomic demands.

Why isoleucine, of all amino acids? The answer might lie in its replaceability. Independent support for isoleucine being the most substitutable of the canonical amino acids comes from a recent effort to build a bacterial cell with a 19-amino acid alphabet, which selected isoleucine for elimination on the basis of low cross-ortholog conservation and computational design tolerance^36^. In that work, 52 essential *Escherichia coli* ribosomal proteins were engineered to replace isoleucine residues with other amino acids. Bacteria with isoleucine-free ribosomes showed minimal fitness phenotypes.

The parasite we used to probe the model, *Plasmodium falciparum*, should be read with survivorship bias in mind. It can grow in adult blood because it induces new permeability pathways (NPPs) that scavenge isoleucine and other nutrients from plasma^23^, making it a survivor of the restriction rather than a naïve subject of it. We view the growth arrest that adult blood still imposes when plasma isoleucine is withdrawn as a footprint of that barrier. Accordingly, it was previously suggested that *Plasmodium* might adjust its growth in response to starvation-induced fluctuations in plasma isoleucine concentrations^24^. Crucially, isoleucine restriction itself is agnostic to any particular parasite. It is not a targeted immune response but a basic property of the meal itself, a food impoverished in one essential amino acid. As such it stands as an obstacle to any blood-borne pathogen that cannot make its own isoleucine. Applying the same framework of amino acid restriction to less-characterized parasites and host proteins could provide a fruitful avenue for future research.

On the host side, our model reframes a classic chapter of human genetics. Malaria has been among the strongest selective forces in recent human evolution, leaving fingerprints across our genome and historical records^37,38^. The protection that hemoglobin variants confer against malaria is the basis of the long-standing malaria hypothesis for the persistence of thalassemias, sickle-cell trait, and other hemoglobinopathies^37,39^. These variants have been read as a set of pathogenic states maintained by parasite pressure. We suggest that the same pressure might have reached the baseline state itself. The isoleucine-free composition of ordinary adult hemoglobin may be a protective feature in its own right, not only through hemoglobin’s pathogenic variants. What functional cost, if any, the exclusion of isoleucine imposes on hemoglobin remains to be understood.

A further implication concerns fetal hemoglobin itself. That *Plasmodium falciparum* escapes isoleucine-withdrawal dormancy on cord blood raises the possibility that fetal hemoglobin, by supplying isoleucine from within the cell, is a generally more permissive substrate for blood parasites under physiological conditions. Our experiments cannot establish this directly and further investigation is required. The question nonetheless has present-day relevance. Exagamglogene autotemcel (exa-cel; marketed as Casgevy), the first approved CRISPR-based therapy, treats sickle cell disease and transfusion-dependent beta-thalassemia by disrupting the erythroid enhancer of BCL11A in a patient’s own hematopoietic stem cells, relieving repression of gamma globin and durably reactivating fetal hemoglobin in the adult red blood cells it produces^40,41^. Such patients therefore carry, in adult circulation, hemoglobin that contains isoleucine. If fetal hemoglobin does relieve the isoleucine restriction we describe, its therapeutic re-expression could in principle lift one of the barriers that ordinarily constrains blood-stage growth of the malaria parasite. This possibility warrants attention given that these therapies are expected to be increasingly deployed in the malaria-endemic regions where sickle cell disease is most common^42^.

Finally, our observations point to the fact that protein composition can be used as a weapon against pathogens. The idea that hosts defend themselves by controlling nutrient availability is not new. Animals withhold trace metals such as iron, zinc, and manganese from invading microbes to starve them^43,44^. What we describe extends this logic from freely exchangeable metals to the fixed composition of a protein itself. Alongside the nutritional, metabolic, and translational pressures already known to shape amino acid usage, we propose that the amino acid composition of an abundant protein functions as a built-in defense against pathogens that would feed on it. Reading protein sequences in this light may reveal further cases in which the code has been shaped not only by what a protein must do, but by what it must deny.

## LIMITATIONS OF THE STUDY

Our study aims to frame molecular patterns in mosquito proteomes and mammalian blood in the light of the hundreds of millions of years of interaction that tie them together. Given the scope and ancient origin of the conflict, it is impossible to establish direct causality. We observe that blood-feeding mosquitoes encode less isoleucine than their relatives that stopped feeding on blood and that this mirrors the isoleucine-poor composition of adult mammalian blood. However, such a signal cannot on its own show that blood feeding drove the shift. Similarly, it is impossible to establish that blood parasite pressure is what drove the removal of isoleucine from adult hemoglobin in mammals. The proteome-wide mosquito inference is further constrained by the structure of the phylogeny. Only three lineages have independently abandoned blood feeding, and because diet, phylogenetic position, and mutational bias covary across these clades, our power to separate the effect of diet from lineage-specific traits is inherently limited. Our readout of amino acid usage is genetic rather than proteomic, inferred from protein-coding sequence and, for the induction analysis in *Aedes aegypti*, from transcriptionally-weighted coding sequence. This captures what genomes encode rather than the proteins actually made. This proxy might be conservative, since isoleucine can in some contexts be functionally substituted by valine without a change to the underlying DNA or RNA sequence^45^.

In our *Plasmodium falciparum* experiments, the comparison of cord blood red blood cells and adult red blood cells introduces potential confounders. Cord blood and adult red blood cells differ in parameters other than their fetal hemoglobin content, which might also affect parasite growth. Additionally, the comparison is not isogenic, and cord blood and adult red blood cell donors differ in their genetic background. Accordingly, we observe variation in growth rate across the different donors, apparent, for example, as slower growth in Adult Blood Donor 2 compared to Adult Blood Donors 1 and 3. These differences might reflect the impact of host genetic variation on *Plasmodium falciparum* proliferation^46^. Our *Plasmodium falciparum* culture medium contains AlbuMAX II, which includes bovine albumin and other blood-derived proteins that may provide the parasite with a limited source of isoleucine, even under the 0 µM extracellular isoleucine condition.

## ACKNOWLEDGMENTS

We thank members of the Vosshall Lab for discussion and comments on the study. We thank Alison Barton, David Reich, and David Zeevi for helpful discussions; Alexandra E. DeFoe for assistance in figure design; and the High-Performance Computing Resource Center at the Rockefeller University for IT support.

## AUTHOR CONTRIBUTIONS

L.H.-Z. designed and performed all computational, genomic, and evolutionary analyses; M.K. performed the *Plasmodium falciparum* culture experiments under the supervision of M.T.D., with statistical analysis and visualization carried out together with L.H.-Z.; L.H.-Z. and L.B.V. conceived the study, designed the figures, and wrote the paper, with input from all authors.

## EXTENDED DATA FIGURES

**Extended Data Figure 1.**
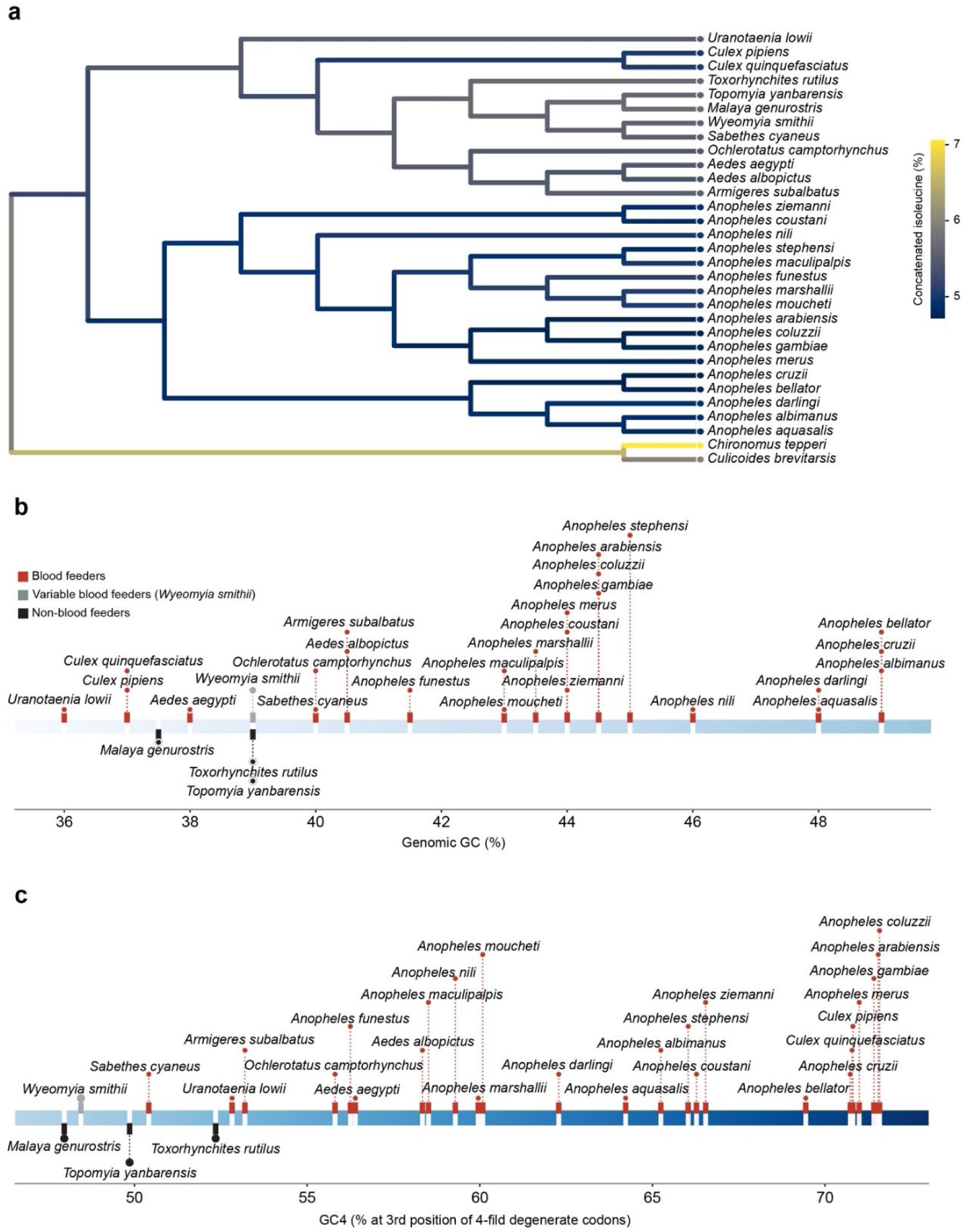
Isoleucine levels, nucleotide composition, and mutational bias across mosquito species. **a,** Species tree of the 29 mosquitoes analyzed together with two dipteran outgroups (*Chironomus tepperi* and *Culicoides brevitarsis*), with branches and tips colored by concatenated-proteome isoleucine content (% of standard residues; scale at right). **b,** Genome-wide GC content (%) for each species, colored by feeding phenotype (blood feeders, red; variable blood feeder *Wyeomyia smithii*, gray; non-blood feeders, black). **c,** Mutational bias, measured as GC4 (GC content at the third position of four-fold degenerate codons; %), colored as in b.

**Extended Data Figure 2.**
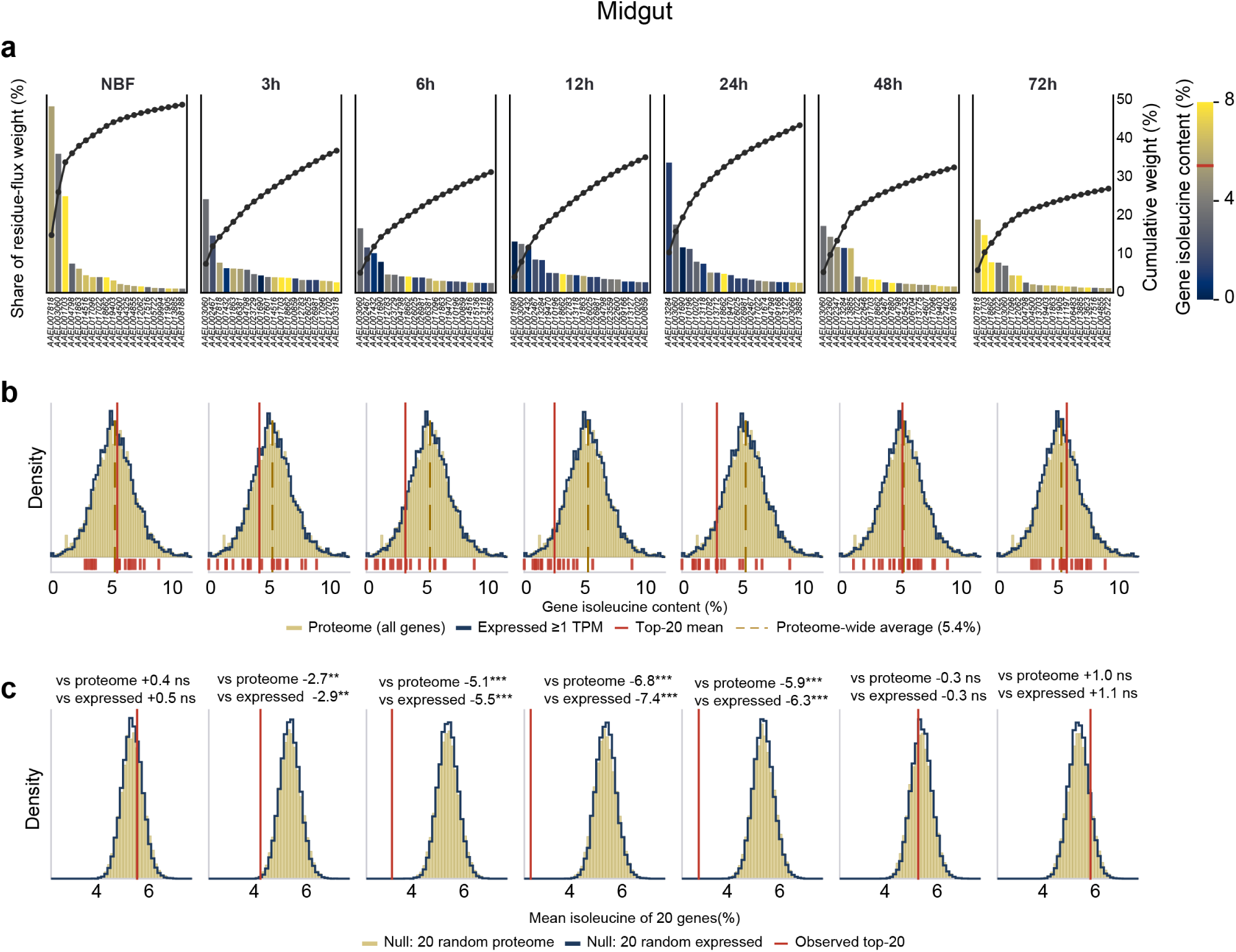
Transcriptionally-weighted usage analysis of the *Aedes aegypti* midgut across a blood-meal time course. **a,** The 20 transcripts contributing most to residue-flux weight at each time point (non-blood-fed, NBF, and 3, 6, 12, 24, 48 and 72 h). Bars show each transcript’s share of total residue-flux weight (left axis) and are colored by gene isoleucine content (% residues, color scale at right; red tick marks the proteome-wide average, 5.4%); Overlaid line shows cumulative weight (right axis). Gene names (AAEL identifiers) are presented below**. b,** Distribution of gene isoleucine content for all proteome genes (tan) and for genes expressed ≥1 TPM in the midgut (blue outline). Red line represents mean isoleucine of the top-20 residue-flux transcripts; dashed line represents proteome-wide average (5.4%); red marks beneath the distribution indicate isoleucine levels in each of the top-20 genes. **c,** Bootstrap test (20,000 iterations) comparing the observed top-20 mean isoleucine (red line) with null distributions of 20 genes drawn at random from the whole proteome (tan) or from genes expressed (>1 TPM) at that time point (blue). Z-score and two-sided empirical significance are given above each panel. The top-20 set is significantly isoleucine-depleted at 3-24 h (peaking at 12–24 h) and returns to baseline by 48–72 h. (**P < 0.01, ***P < 0.001; ns, not significant.)

**Extended Data Figure 3.**
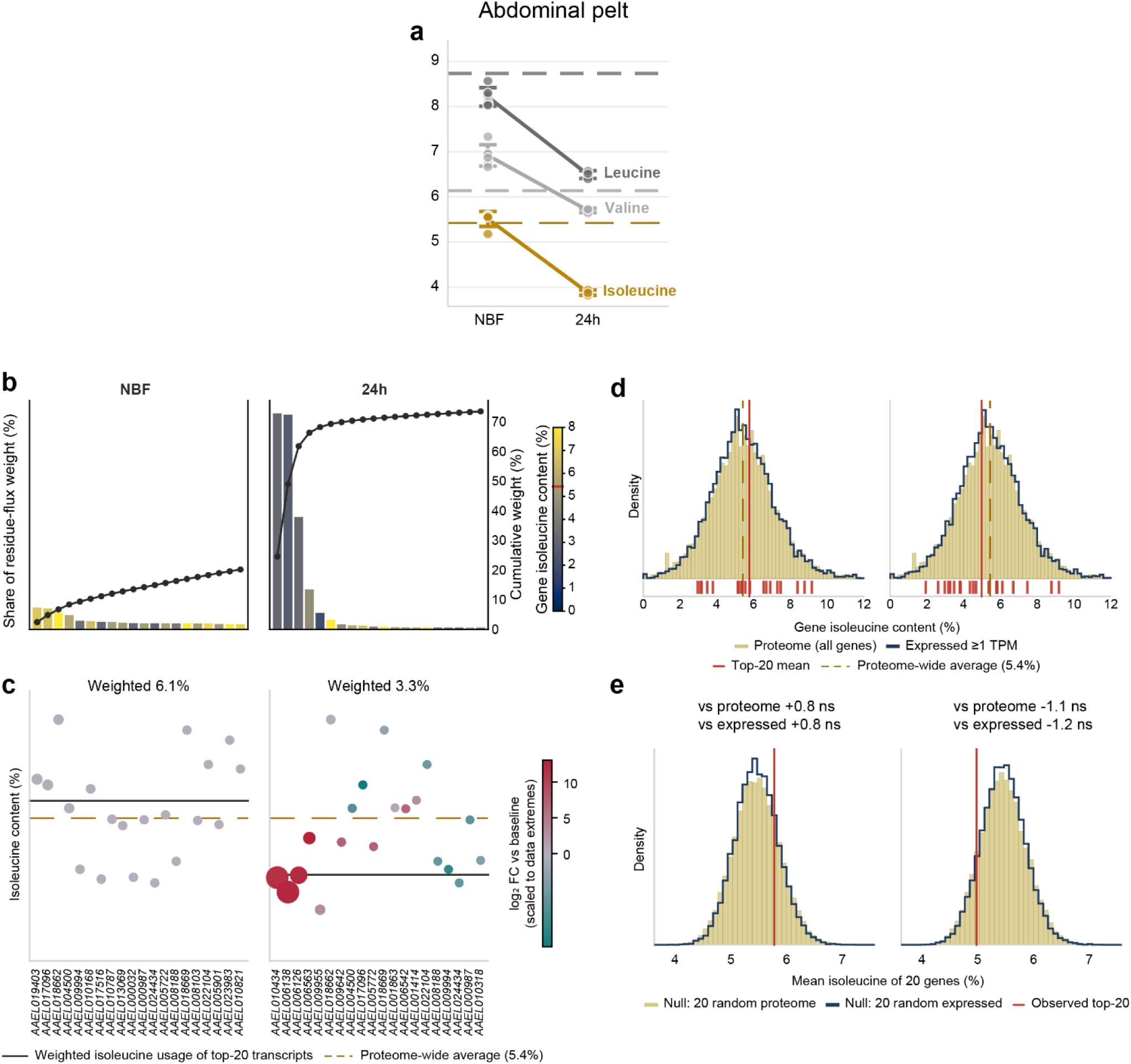
Transcriptionally-weighted usage analysis of the *Aedes aegypti* abdominal pelt before and after a blood meal. *Aedes aegypti* abdominal pelt (fat body), non-blood-fed (NBF) versus 24 h after a blood meal. **a,** Transcriptionally-weighted usage of isoleucine (gold), valine and leucine (gray) at non-blood-fed (NBF) versus 24 h after a blood meal; points are biological replicates and dashed lines the proteome-wide averages. **b,** The 20 transcripts contributing most to residue-flux weight at each time point. Bars show each transcript’s share of total residue-flux weight (left axis) and are colored by gene isoleucine content (% residues, color scale at right; red tick marks the proteome-wide average, 5.4%). Overlaid line shows cumulative weight (right axis). **c,** Isoleucine content of the top-20 transcripts (points sized by residue-flux share and colored by log_2_ fold-change versus non-blood fed samples); solid line, weighted isoleucine of the top-20 (6.1% at NBF, 3.3% at 24h); dashed line, proteome-wide average (5.4%). Gene names (AAEL identifiers) are presented below. **d,** Isoleucine-content distributions at NBF and 24 h, as in Extended Data Fig. 2b. **e,** Bootstrap test as in Extended Data Fig. 2c.

**Extended Data Figure 4.**
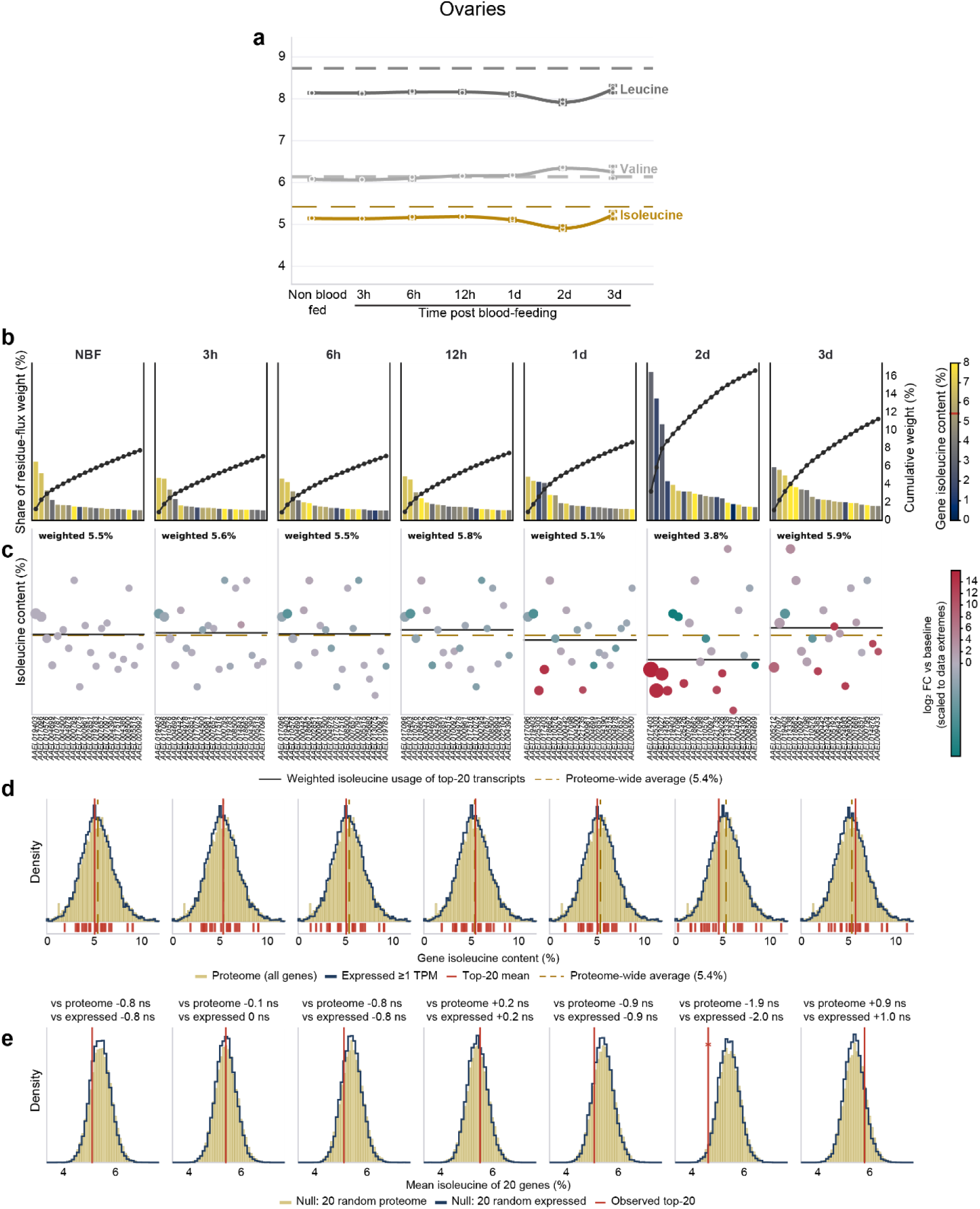
Transcriptionally-weighted usage analysis of the *Aedes aegypti* ovaries across a blood-meal and reproductive time course. *Aedes aegypti* ovaries across a blood-meal time course (NBF and 3, 6, 12 h, 1, 2 and 3 d). **a,** Transcriptionally-weighted usage of isoleucine (gold), valine and leucine (gray) over time; dashed lines, proteome-wide averages. **b,** The 20 transcripts contributing most to residue-flux weight per time point, as in Extended Data Fig. 2a. **c,** Isoleucine content of the top-20 transcripts, sized by residue-flux share and colored by log_2_ fold-change versus non-blood fed samples. Gene names (AAEL identifiers) are presented below and are shared with b. **d,** Isoleucine-content distributions per time point, as in Extended Data Fig. 2b. **e,** Bootstrap test per time point, as in Extended Data Fig. 2c.

**Extended Data Figure 5.**
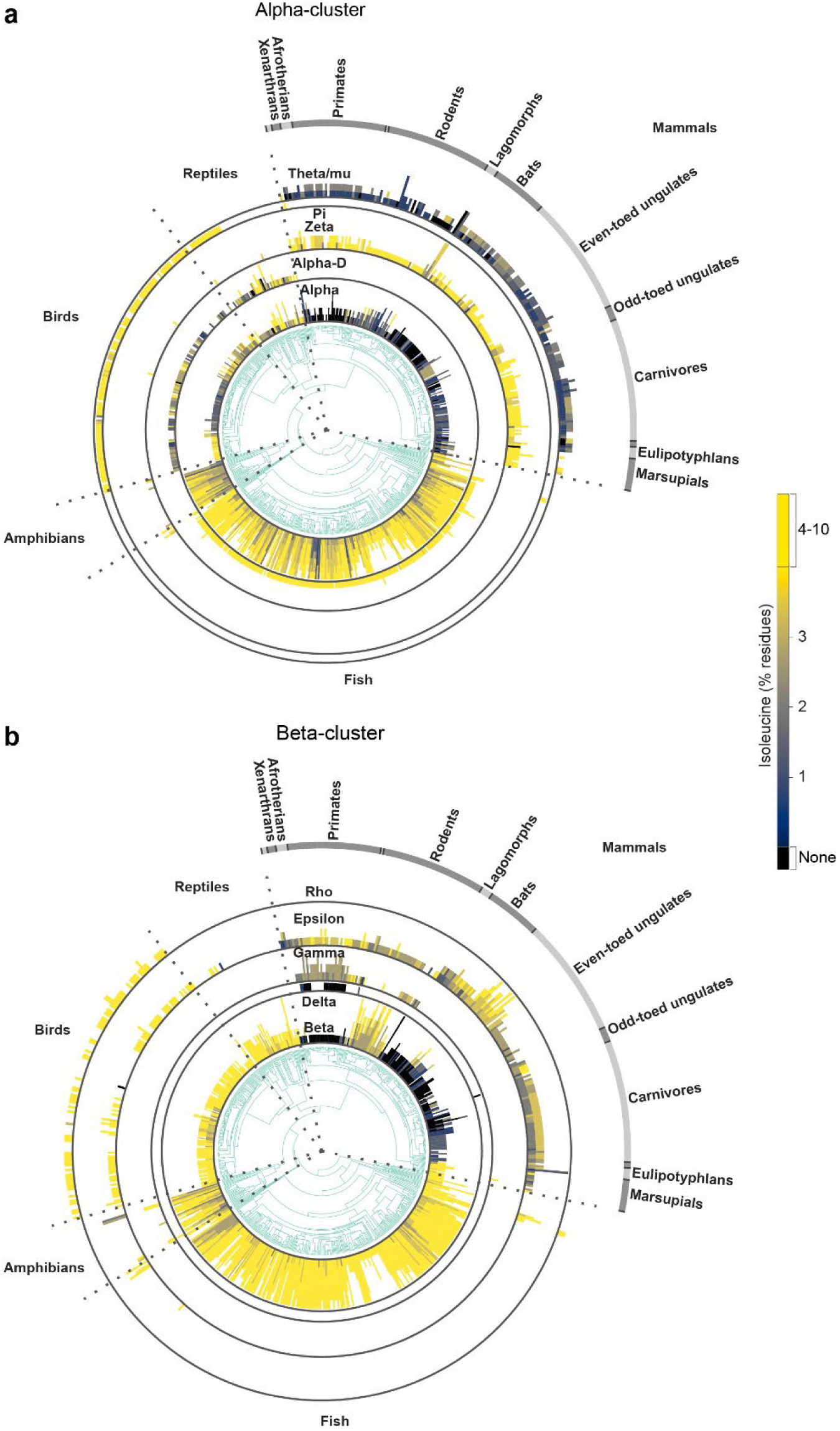
Isoleucine content of hemoglobin subunit types across vertebrates. Hemoglobin proteins from 707 vertebrate genomes (658 species plotted on tree) placed on the vertebrate species tree and split into the alpha and beta clusters. Each bar is one gene, colored by isoleucine content (black, none; blue-to-yellow gradient up to 4-10% of residues; scale at right). Concentric rings correspond to inferred subunit type; outer arc labels vertebrate and mammalian groups. **a,** Alpha-cluster subunit types (alpha, alpha-D, zeta, pi, theta/mu). **b,** Beta-cluster subunit types (beta, delta, gamma, epsilon, rho).

**Extended Data Figure 6.**
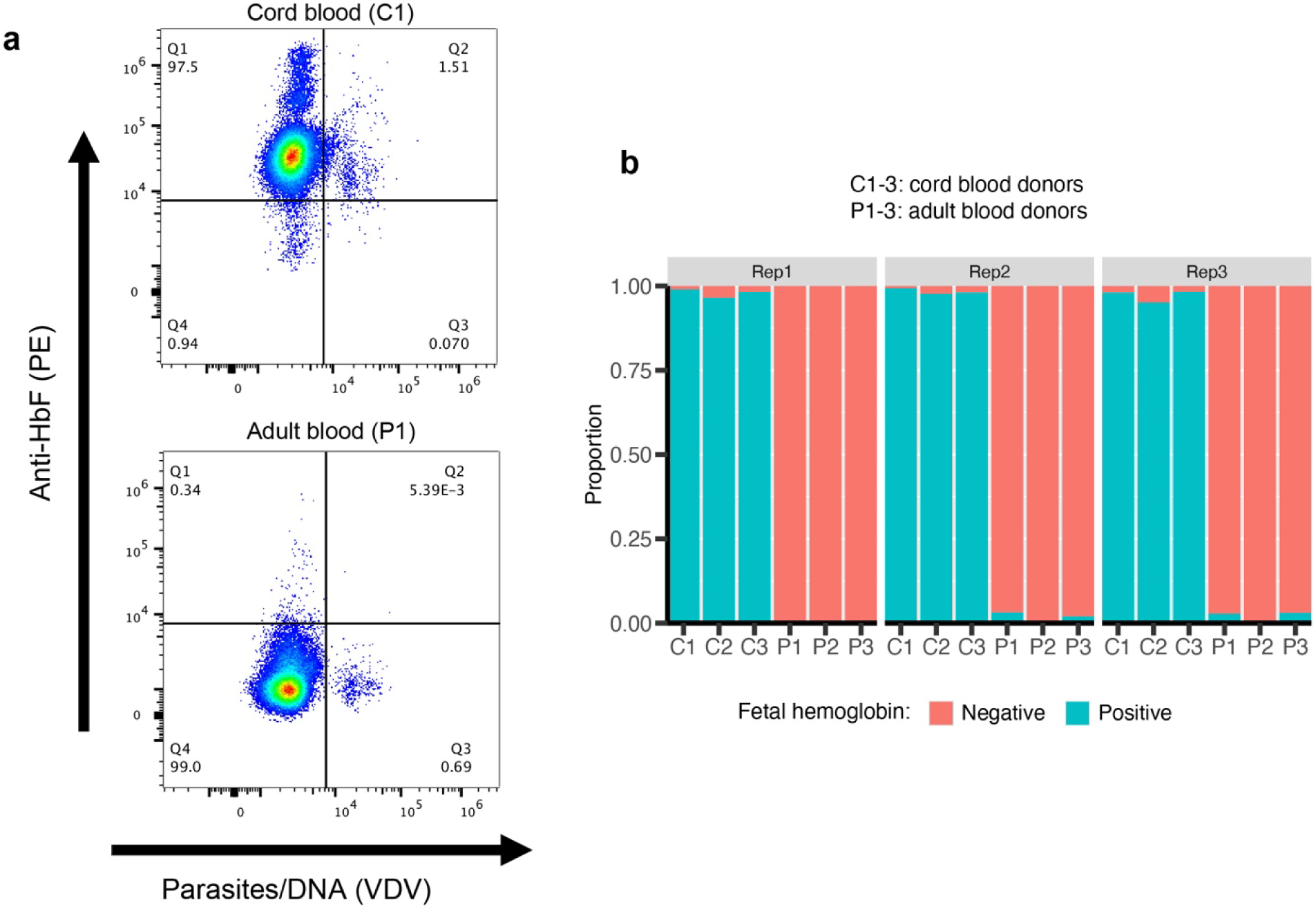
Fetal hemoglobin content of the cord and adult red blood cells used in *Plasmodium falciparum* growth assays. **a**, Representative flow cytometry plots of red blood cells from one cord blood donor (C1, top) and one adult blood donor (P1, bottom), collected at assay setup, fixed, permeabilized and stained with anti-HbF–PE (y axis) and the DNA dye Vybrant DyeCycle Violet (VDV, x axis), which marks parasite DNA. Numbers give the percentage of single cells in each quadrant. Cord blood cells are almost uniformly HbF positive (Q1 + Q2), whereas adult blood cells are HbF negative (Q3 + Q4). **b**, Proportion of HbF-positive (teal) and HbF-negative (salmon) red blood cells for each of the three cord (C1-C3) and three adult (P1-P3) blood donors, shown for each of three biological replicates (Rep1-Rep3).

**Extended Data Figure 7.**
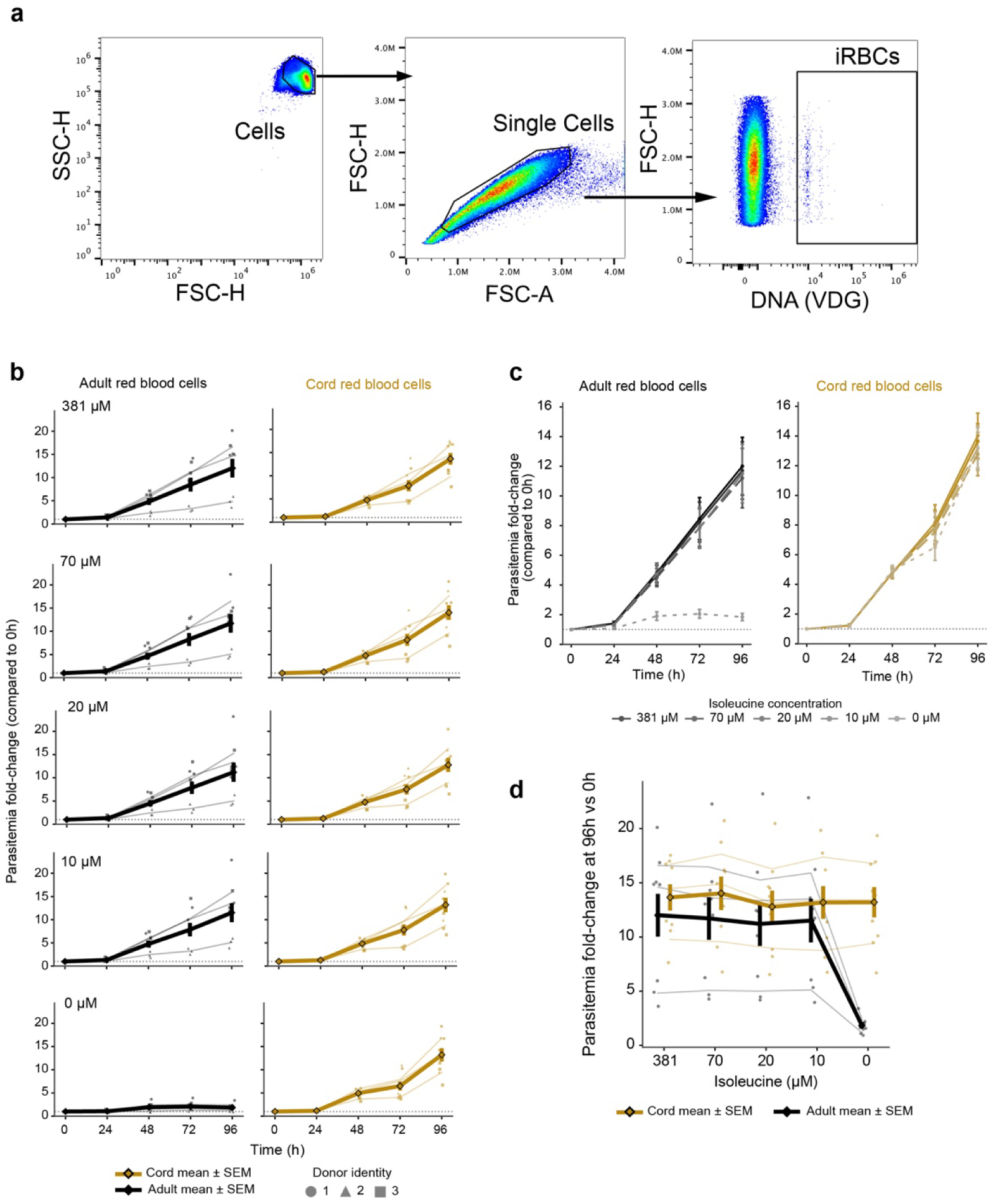
Gating strategy and isoleucine dose-response of *Plasmodium falciparum* growth in adult versus cord red blood cells. **a**, Flow cytometry gating strategy used to quantify parasitemia. Cells were gated on SSC-H versus FSC-H (left), single cells on FSC-H versus FSC-A (middle), and infected red blood cells (iRBCs) identified as Vybrant DyeCycle Green (DNA, VDG)-positive events (right). **b**, Parasitemia fold-change relative to 0 h over 96 h for *Plasmodium falciparum* cultured in adult (black, left) or cord (fetal-hemoglobin-containing; gold, right) red blood cells, across a range of extracellular isoleucine concentrations (top to bottom: 381, 70, 20, 10 and 0 µM). Individual points denote n = 3 independent blood donors per blood category, with donor identity indicated by symbol shape (donor 1, circle; donor 2, triangle; donor 3, square); thin lines show per-donor means across biological replicates. Thick lines denote the group mean and error bars the standard error of the mean across donors. The dotted horizontal line marks no change relative to t0 (fold change = 1). **c**, Overlay of all isoleucine concentrations for adult (left) and cord (right) red blood cells, showing parasitemia fold change over 96 h. Line shade indicates isoleucine concentration (dark to light: 381 to 0 µM); the no isoleucine (0 µM) condition is shown as a dashed line. Thick lines and error bars denote the group mean ± s.e.m. across donors (n = 3). **d**, Endpoint (96 h) parasitemia fold change as a function of extracellular isoleucine concentration (381 to 0 µM), for adult (black) and cord (gold) red blood cells. Individual points denote per-donor values colored by blood category; thin lines show per-donor trajectories across concentrations; thick lines and error bars denote the group mean ± s.e.m. across donors (n = 3).

**Extended Data Figure 8.**
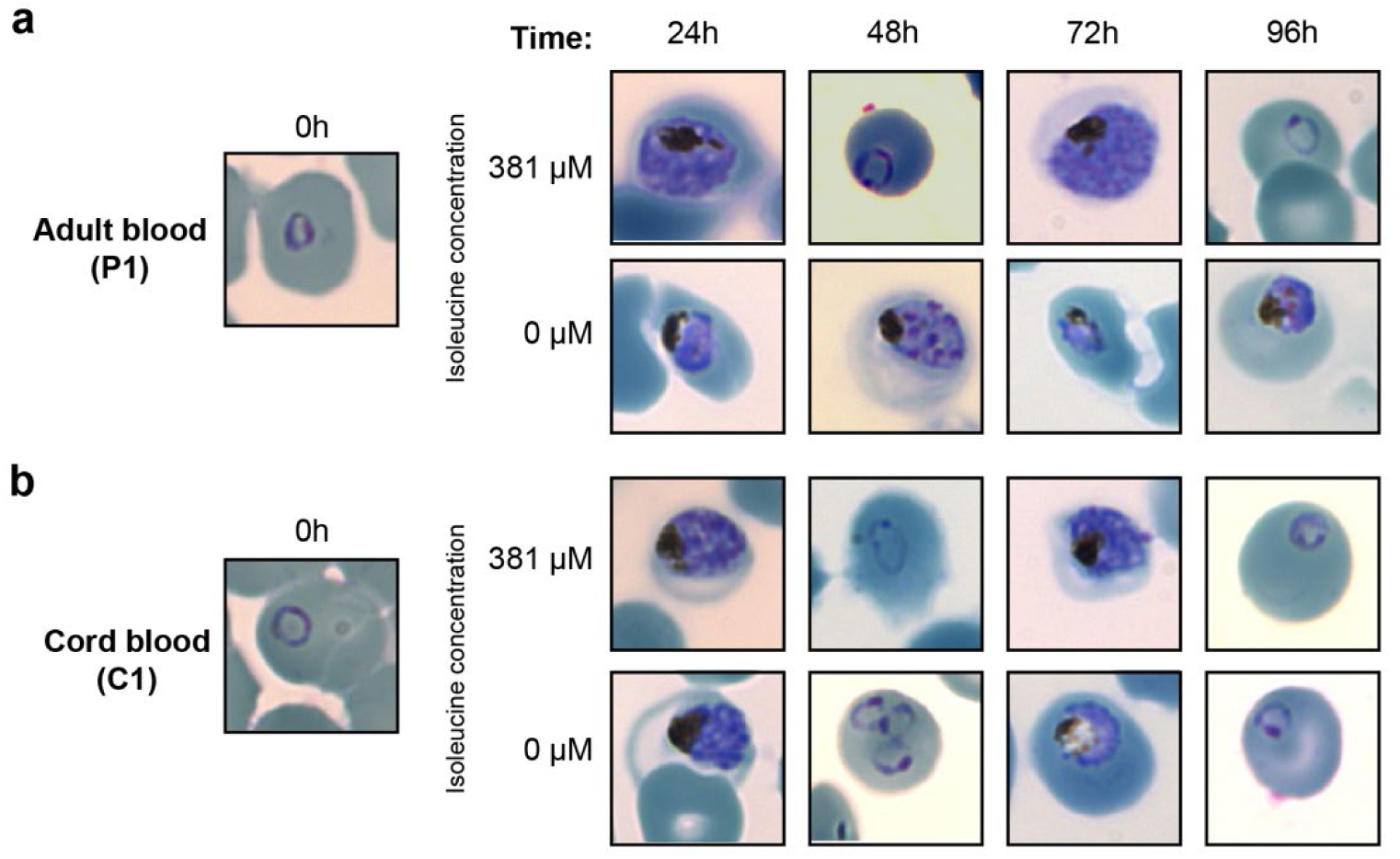
Morphology of *Plasmodium falciparum* in adult and cord red blood cells with and without extracellular isoleucine. **a,** Representative micrographs of Hemacolor-stained blood smears of *Plasmodium falciparum* cultured in adult red blood cells (donor P1) at assay setup (0 h, left) and at 24, 48, 72 and 96 h in medium containing 381 µM (top) or 0 µM (bottom) extracellular isoleucine. **b,** As in a, for parasites cultured in cord (fetal-hemoglobin-containing) red blood cells (donor C1). Smears are from one representative donor per blood category; parasitemia and stage composition quantified across all three donors per category are shown in Fig. 4e.

## MATERIALS AND METHODS

### Mosquito phylogeny and host preference

The estimated age of the mosquito family was retrieved from Soghigian et al.^3^. The visualization of the phylogenetic arrangement of the 29 mosquito species in this study is based on the species tree output of orthology analysis (OrthoFinder, see below).

Mosquito blood-feeding host preference was assigned for each of the 29 mosquito species analyzed in this study. The primary source was the per-species blood-meal count table of Soghigian et al.^3^ (counts mirrored from the authors’ Culicitree repository, supplementary_data_4 / MOESM8), which provided molecular blood-meal records for 22 of the 29 species. For each species, recorded meals were tallied across four host classes: amphibians, reptiles, birds and mammals (humans included within mammals). Each class was scored as a major host (≥20% of recorded meals), an occasional host (1-20%), or absent (<1% or no records).

Seven species absent from Soghigian et al.^3^ were resolved by targeted literature review: three obligate non-blood-feeders (*Malaya genurostris, Topomyia yanbarensis* and *Toxorhynchites rutilus*) were coded as non-feeders, the geographically variable feeder *Wyeomyia smithii* was coded separately, and three further blood-feeders (*Anopheles merus*, *Anopheles bellator*, and *Sabethes cyaneus*) were assigned from species-level literature. A per-species confidence flag was recorded from the underlying count (high ≥100, moderate 30-99, low <30 meals). The full per-species assignments, host-class counts and proportions, primary references and confidence flags are compiled in the accompanying workbook and provided in the Extended Data^14^.

### Analyzing amino acid usage in mosquito proteomes

#### Genome set, isoform collapse and transposon removal

Protein-coding sequences for the 29 mosquito species and the outgroup taxa used in the cross-species comparison were retrieved from NCBI RefSeq. For each assembly, the protein FASTA and the feature table were downloaded directly from the RefSeq FTP tree by accession. Isoforms were collapsed to one representative per gene by parsing the feature table and retaining, for every gene, the longest protein isoform among its XP_ records. The corresponding protein sequences were extracted from the RefSeq protein FASTA.

Transposable-element (TE)-derived proteins were then filtered out. Each collapsed proteome was annotated with InterProScan (applications Pfam and SUPERFAMILY). Proteins were flagged as candidate TEs when a domain description matched a fixed keyword list (case-insensitive substrings: transpos, RT_L, RT_N, reverse transcriptase, integrase, Gypsy, DUF5641, DUF1759, retro), scanned against the InterProScan signature- and InterPro-description fields.

#### Amino acid usage measures

For each clean proteome, amino acid usage was quantified in two ways.

(1) Concatenated-proteome usage: all proteins of a species were pooled and each of the 20 standard amino acids scored as its total count divided by the total number of standard residues (non-standard characters ignored) X 100.
(2) Median per-protein usage: each protein’s composition (percentage of residues) was computed and the median taken across proteins per amino acid. Isoleucine content is reported as the percentage of isoleucine among standard residues.

Each measure was computed over three protein sets: (a) the full protein-coding repertoire; (b) proteins present in all species, regardless of copy-number; and (c) single-copy (1-to-1) orthologs shared across all species. (b) and (c) defined from the orthology inference below.

#### Orthology inference

Orthogroups and single-copy orthologs were inferred with OrthoFinder^47^ (v3.1.3) from the clean, longest-isoform proteomes, run in multiple-sequence-alignment mode (orthofinder -f <COLLAPSED_FASTAS>/ -t 32 -M msa). In addition to the 29 mosquito species, two non-mosquito Diptera were included as outgroups – *Culicoides brevitarsis* (a biting midge, *Ceratopogonidae*) *and Chironomus tepperi* (a non-biting midge, *Chironomidae*). Hierarchical orthogroups (HOGs) were taken at node N2, the internal node corresponding to the common ancestor of the 29 mosquitoes (i.e. excluding the two outgroups), so that the “present in all species”, single-copy (1-to-1) ortholog sets, and the ortholog mappings used in the cross-species comparisons are defined within the mosquito clade.

### Phylogenetic test of isoleucine-encoding residues across mosquito species

#### Phylogenetic framework

Species relationships and divergence times were taken from the dated maximum-clade-credibility mosquito phylogeny of Soghigian et al.^3^ (mcc_alltribes.tre, 3,567 tips), pruned to the 29 focal species. *Ochlerotatus camptorhynchus* is represented in that phylogeny as *Aedes camptorhynchus.* Phylogenetic covariance between species was defined as the root-to-most-recent-common-ancestor path length for each pair.

#### Test of isoleucine usage against feeding state

Isoleucine usage, expressed as the percentage of isoleucine among standard residues in the concatenated proteome, was regressed on feeding state by phylogenetic generalized least squares (PGLS). Tests were performed on the full protein-coding repertoire, pre-specified as the primary gene set. Usage over proteins present in all species and over single-copy orthologs is shown descriptively (Fig. 1e, Extended Data^14^). Two groupings of feeding state were analyzed, defined a priori: the three lineages that have abandoned blood feeding (*Toxorhynchites rutilus*, *Topomyia yanbarensis*, *Malaya genurostris*) versus all other species, and those three together with the variable blood feeder *Wyeomyia smithii* versus all other species. All parametric p values are two-tailed.

Pagel’s λ was estimated jointly with the regression coefficients by maximum likelihood over the interval [0, 1], with a 95% confidence interval from profile likelihood (a decrease of 1.92 log-likelihood units). λ was intermediate (0.88-0.89, 95% CI 0.63-0.97) and differed significantly from both a Brownian model (λ = 1; likelihood ratio 14.2-14.8, p ≤ 1.7 × 10⁻⁴) and phylogenetic independence (λ = 0; likelihood ratio 29.4-32.8, p ≤ 6.0 × 10⁻⁸).

#### Non-parametric confirmation and sensitivity analyses

We additionally performed a phylogenetic simulation test (phylogenetic ANOVA^48^): under the fitted intercept-only model, 10,000 trait vectors were simulated along the phylogeny using the estimated λ and residual variance, the full model was refitted to each with λ re-estimated by maximum likelihood, and the empirical p value was taken as the proportion of simulated *t* statistics at least as extreme, in the predicted direction, as the observed value. As an assumption-free check we also enumerated exhaustively all possible assignments of the observed group sizes to the 29 tips (3,654 and 23,751 labelings for the two groupings respectively) and recomputed the test statistic for each.

Results were insensitive to the time calibration: replacing branch lengths with Grafen’s transform of the same topology, which discards divergence times, gave p = 0.046 and 0.035 for the two groupings. Model choice was assessed by comparing Brownian, Pagel’s λ and single-optimum Ornstein-Uhlenbeck models of trait evolution. The λ model was best supported (ΔAIC ≥ 9.7), and the estimated effect of feeding state was of the same sign and comparable magnitude under each.

#### Genomic GC and GC4 analysis

Two measures of nucleotide composition were compared against blood-feeding state across the 29 mosquito species. Genome-wide GC content was taken from the NCBI assembly statistics for each species. GC4 (GC content at the third position of four-fold degenerate codons, a relatively neutral proxy for mutational bias^15^) was computed from each species’ filtered coding sequences. Eight four-fold degenerate codon families were examined: Ala, Gly, Pro, Thr and Val, plus the four-fold blocks of Leu [CTN], Ser [TCN] and Arg [CGN]. The fraction of third-codon positions that are G or C was measured. Coding sequences whose length was not a multiple of three were skipped. Each species was placed at its genomic-GC and its GC4 value and colored by feeding strategy (blood feeders versus non-blood feeders).

We additionally refitted the PGLS model with GC4 as an additional covariate, with λ re-estimated. GC4 predicted isoleucine usage strongly (-0.025 percentage points of isoleucine per percentage point of GC4), but the coefficient for feeding state remained negative in both groupings (β = -0.11, p = 0.011 for the three obligate non-blood feeders; β = -0.08, p = 0.075 including *Wyeomyia smithii*). With GC4 included, Brownian, λ and Ornstein-Uhlenbeck models converged (λ -> 1, α -> 0) and gave essentially identical estimates of the effect of feeding state (β = -0.111 to -0.110).

### Transcriptionally-weighted amino acid usage

#### RNA-sequencing datasets and processing

Blood-meal transcriptional responses were analyzed for three *Aedes aegypti* tissues using published RNA-seq datasets: midgut across a blood-meal time course^10^ (non-blood-fed baseline and 3, 6, 12, 24, 48 and 72 h post-blood-meal), ovary^49^ (non-blood-fed baseline and 3 h, 6 h, 12 h, 1 d, 2 d and 3 d, the last eggs-retained), and abdominal pelt/fat body^50^ (pooled controls as baseline and 24 h blood-fed). The blood-meal RNA-seq datasets were processed uniformly through a single pipeline: adapter and quality trimming with Cutadapt^51^, transcript quantification with kallisto^52^, and gene-level summarization with tximport^53^, as described below. Each dataset was processed independently.

Read quality was assessed with FastQC^54^. When needed, paired-end reads were then trimmed with Cutadapt^51^ (v5.2) using these Illumina TruSeq adapter sequences:

AGATCGGAAGAGCACACGTCTGAACTCCAGTCA (read 1, -a)

AGATCGGAAGAGCGTCGTGTAGGGAAAGAGTGT (read 2, -A).

We used a quality cutoff of 20 (-q 20) and a minimum retained read length of 36 nt (-m 36). Trimmed reads were quantified against the *Aedes aegypti* reference transcriptome (VectorBase AaegLVP, build 58 “Jov19” annotation^55^) with kallisto^52^ (v0.52.0) in paired-end mode with 100 bootstrap replicates (-b 100). For samples sequenced across more than one lane, all lane-level FASTQ pairs belonging to a biological sample were supplied to a single kallisto run, so that lanes were pooled at the quantification step. Transcript-level abundances were summarized to gene level with tximport, using a transcript-to-gene map derived from the matching GFF annotation. Per-gene transcripts-per-million (TPM) values were used for all downstream analyses.

#### Transcriptionally-weighted usage of amino acids

To measure the translational output and amino acid usage of a tissue, we weighted each protein’s amino acid composition by its expression and its length. For each gene, we defined a residue-flux weight equal to its expression times its protein length (w_g = TPM_g × L_g, where L_g is the protein length in standard residues). The weighted usage of an amino acid at a given condition is the total number of that amino acid’s residues across all proteins, weighted by expression, divided by the total number of weighted residues:

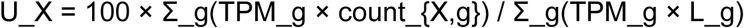

Weighted usage was computed separately for each replicate and summarized as the mean ± s.d. across replicates at each timepoint, for isoleucine, valine and leucine. As a reference, the dashed line marks the proteome-wide usage of each amino acid (∼5.4% for isoleucine).

#### Top-20 residue-flux transcripts and bootstrap

At each timepoint, per-gene mean TPM across the condition’s replicates gave w = TPM × L, and the 20 genes with the largest weight were taken as top contributors. Each contributor’s isoleucine content, its share of total residue-flux weight, and its log_2_ fold-change versus the non-blood-fed baseline were reported. To test whether the top-20 set is isoleucine-depleted as a group, the mean isoleucine of the top-20 was compared with the mean of 20 genes drawn at random (20,000 iterations) from (i) the whole proteome and (ii) genes expressed (>1 TPM) in the tissue; a z-score and two-sided empirical p-value were computed for each null.

The per-gene composition table used for weighted usage is the full VectorBase (build 68) *Aedes aegypti* protein-coding set (14,718 genes, longest isoform per gene), matched to the TPM matrices by AAEL. Where a downstream step was restricted to RefSeq-orthologous genes for cross-species comparison, the VectorBase-only genes drop out. Scope was kept consistent within a single figure.

### Isoleucine in domain-sharing genes and orthologs

#### Focal blood-meal-expressed gene selection

Blood-feeding expressed midgut genes were selected from the residue-flux ranking. Per timepoint, each gene’s weight w = TPM × L was normalized to a share of total flux. A combined rank summed each gene’s weight-share across the midgut window (6, 12 and 24 h) and ranked genes descending. The top-20 combined-rank genes were taken as focal. Gene IDs are provided in Extended Data Figs. 2-4 and Extended Data^14^.

#### Domain-family comparison

Every *Aedes aegypti* protein was annotated with its InterProScan domains; for each focal gene, its most specific shared domain (the least-common domain it carries that is shared by ≥2 genes genome-wide) was taken, and all *Aedes* genes carrying that domain were pooled as the family. Isoleucine content per gene was the mean over its isoforms. Focal genes sharing a domain were collapsed into one family; each family’s isoleucine distribution was compared with the isoleucine content of its focal, blood-feeding-expressed member(s), with family and genomic medians drawn as references.

#### Cross-species orthologs of blood-feeding-expressed genes

Orthologs of the focal blood-feeding-expressed midgut genes were compared across species using OrthoFinder hierarchical orthogroups at the N2 node. Each focal AAEL gene was mapped to its *Aedes* protein and then to its N2 HOG; all HOG member proteins in each species were retrieved and their isoleucine content computed. Paralogs sharing a HOG were kept as separate values. Species were grouped by feeding strategy: blood feeders, the three non-blood-feeders (*Topomyia yanbarensis, Malaya genurostris, Toxorhynchites rutilus*), and the frog-blood-feeding *Uranotaenia lowii*. Genes mapping to no N2 HOG were omitted (in the combined top-20 list, cytochrome c oxidase *AAEL018662* and a mucin-like peritrophin AAEL004798).

### Vertebrate hemoglobin analysis

#### Vertebrate globin gene set

Protein FASTA files for 707 vertebrate species were retrieved from NCBI RefSeq (full species and accession list in the Extended Data^14^). Globin-domain proteins were identified by searching the Pfam Globin profile HMM (PF00042) against each proteome with hmmsearch, using Pfam’s curated gathering threshold for family membership. Two filters were then applied: hits whose RefSeq name contained “LOW QUALITY PROTEIN” were dropped and logged, and sequences longer than 170 aa were removed, excluding large multidomain globin-containing proteins whose PF00042 hit is only a fragment (androglobin, ∼1,600 aa, and fusions up to ∼3,000 aa) while retaining the canonical respiratory globins: hemoglobin α/β (∼141–146 aa), myoglobin (∼154 aa) and neuroglobin (∼151 aa). This yielded 9,639 globin proteins across 707 species, each labeled Genus_species_accession and accompanied by a table of species, accession and RefSeq description.

#### Globin gene tree

All retained globins were concatenated into one FASTA, aligned with MAFFT^56^ (--auto --reorder), and a gene tree built with FastTree under the LG model with Gamma rate variation (-lg -gamma); node labels are SH-like local supports. The tree was used to quality-control subunit labels (below) and for the auxiliary ring figures. Alignment and tree are provided in the Extended Data^14^.

#### Isoleucine content and copy selection

Isoleucine content of each protein was computed as the percentage of isoleucine among its residues; sequences with exactly zero isoleucine were plotted in black. Valine and leucine were computed identically for the corresponding panels.

A gene copy was plotted if it passed three cumulative filters: sequences whose RefSeq description contained “partial” were dropped (n = 60); 35 manually curated mammalian entries were removed due to abnormal length, and 16 replaced or reclassified (below); α- or β-cluster copies shorter than 120 aa were dropped, removing truncated hemoglobin fragments (∼1% of Hb copies).

#### Cluster and subunit assignment

Each copy was assigned a cluster and subunit from its normalized RefSeq description (lower-cased, with PREDICTED:, isoform, homeolog and “-like” tags stripped) by keyword in fixed priority order; within the α-cluster ζ, α-D, θ/μ, π, otherwise α; within the β-cluster ε, γ, δ, ρ, otherwise β. Assignment follows globin biology, with π drawn in the α figure and ρ in the β figure.

#### Species backbone and vertebrate class

Figures were drawn on the time-calibrated vertebrate species timetree (TimeTree, retrieved 1 July 2026; ultrametric), with one tip per species rather than one per genus, so that genera with several sequenced species contribute several adjacent tips. Of the 707 species carrying globins, 658 matched a timetree tip; the 49 species with no tip were dropped and are listed in the Extended Data^14^. The tree was pruned to the matched tips and single-child nodes were collapsed with branch lengths summed to preserve the time calibration. Each species was assigned a vertebrate class (Mammals, Birds, Reptiles, Amphibians, and the fish grades merged into Fish) from the NCBI Taxonomy, queried offline through taxoniq. Within Mammals, species were further assigned to one of fourteen groups (Monotremes, Marsupials, Xenarthrans, Afrotherians, Primates, Colugos, Rodents, Lagomorphs, Bats, Eulipotyphlans, Pangolins, Carnivores, Even-toed ungulates and Odd-toed ungulates) from the same taxonomy, and these are drawn as labeled segments within the Mammals arc. Ring height was capped at six copies per subunit so that high-copy groups do not dominate the bars; species exceeding the cap are reported with their true counts in an overflow table in the Extended Data^14^. The final backbone carries 658 tips and 8,796 gene copies.

#### Sequence quality control and corrections

Corrections applied at plot time came from four sources. Mammalian labels were cross-checked against the globin gene tree: for each mammalian tip, the smallest enclosing clade of ≥6 tips was polled for its consensus cluster and subunit, and three genes labeled as α-cluster subunits that fall within the β-cluster were reassigned; 38 within-cluster subtype disagreements were recorded but not applied, as recent duplication and gene conversion make within-cluster orthology unreliable. Loci where the longest-isoform rule had retained an over-long mis-predicted isoform were replaced with the canonical RefSeq isoform. Primate and mammalian α/β entries were additionally curated by hand. Finally, four RefSeq entries of the form “hemoglobin subunit alpha-3/-4” (one in *Pan troglodytes*, two in *Pteronotus mesoamericanus*, one in *Bubalus bubalis*) are supernumerary alpha-cluster loci rather than main adult alpha subunits; they were reassigned to a separate non-canonical-alpha subtype and excluded from the main alpha bin used for Fig. 3a and 3d (they remain visible in the subtype panels of Extended Data Fig. 5). None of the four affected species loses a canonical alpha copy as a result.

#### Outputs and software

Per-copy tables (species, cluster, subunit, residue percentage), overflow tables for species exceeding the copy cap, and the list of excluded genes with reasons are provided in the Extended Data^14^. Analyses used Python 3.12 with Biopython, NumPy, Matplotlib and taxoniq.

### Isoleucine usage across the human proteome

Isoleucine content was computed per human protein as count(“I”)/length × 100, and the distribution across proteins used to place the hemoglobin subunits as outliers. The input was the RefSeq human proteome. Predicted models (XP_ accessions) were excluded; only reviewed RefSeq proteins (NP_ accessions) were kept, restricted to ≥ 120 aa. n = 14,296. The distribution is unimodal and slightly right-skewed: mean 4.38%, median 4.27%, SD 2.01%, range 0–14.97%, with 48 proteins at 0% isoleucine. The 4.38% mean is the human proteome average quoted in the manuscript.

#### Albumin and immunoglobulin G analysis

Isoleucine content of human serum albumin was determined from its UniProt sequence (UniProt P02768). The N-terminal signal peptide and propeptide (residues 1-24; the “prepro” region removed during maturation) were excluded, and isoleucine content was computed for the mature 585-residue chain as the number of isoleucine residues divided by the total number of residues. Mature human serum albumin contains 8 isoleucine residues (1.4%).

Isoleucine content was determined for the constant regions of assembled human immunoglobulin G. Constant-region sequences were obtained from UniProt for the IgG1 heavy chain (γ1; UniProt P01857, secreted isoform, 330 residues, 5 isoleucine residues), the IgG2 heavy chain (γ2; UniProt P01859, 326 residues, 4 isoleucine residues), the kappa light chain constant region (κ; UniProt P01834; 107 residues, 1 isoleucine), and the lambda light chain constant region (λ1; UniProt P0CG04; 106 residues, 1 isoleucine). An assembled immunoglobulin was modeled as a tetramer comprising two identical heavy chains and two identical light chains of a single type (κ or λ). Total constant-region residue counts and isoleucine counts were therefore calculated as twice the per-chain values. For an IgG1/κ molecule this yielded 12 isoleucine residues among 874 constant-region residues (1.37%); for IgG1/λ, 12 among 872 (1.38%); and for IgG2/κ, 10 among 866 (1.15%). Variable-domain (VH and VL) residues were excluded from all calculations, as their sequences vary by clone. Isoleucine frequencies were expressed as a percentage of total constant-region residues.

#### Highly expressed proteins in non-blood tissues

The most abundant proteins of several human solid tissues were compared with the major blood proteins. Per-tissue protein abundances were taken from PaxDb integrated tissue (organ) datasets. Weighted-average abundances in ppm of the tissue proteome (pct_of_total_protein = ppm/10,000) were calculated for four tissues: brain, heart, liver, and lung (PaxDb integrated dataset IDs 3536693818, 440180531, 4032974247, and 3105119500 respectively; dataset publication year 2026). Within each tissue the top-3 proteins were taken after excluding hemoglobin and albumin (to avoid blood-contamination proteins). Each protein was matched to its sequence by Ensembl protein identifier (ENSP; provided directly in the PaxDb string_external_id column) using the Ensembl pep.all FASTA and isoleucine content computed as count(“I”)/length × 100; identifications were cross-checked against an independent gene-symbol-to-RefSeq mapping. For blood, canonical literature values were used for whole-blood/plasma composition and isoleucine content (hemoglobin 80%, serum albumin 12%, IgG 3.5%), since PaxDb has no dedicated blood sample.

### Amino acid usage in the Plasmodium falciparum proteome

The amino acid composition of the *Plasmodium falciparum* proteome (RefSeq GCF_000002765.6) was computed by parsing the parasite protein FASTA and calculating each of the 20 standard amino acids as its count divided by the total number of standard residues; amino acids were ranked by the highest overall percentage to lowest.

### Plasmodium falciparum growth assays

Cord blood and adult blood samples were obtained from HumanCellsBio. Cord blood donors were C1 (W243426001091U), C2 (W243426001765F) and C3 (W243426009633). Adult (non-cord) blood control donors were P1 (HH049), P2 (HH061) and P3 (AL003). *Plasmodium falciparum* 3D7 parasites were cultured in the corresponding donor RBCs in complete parasite medium (RPMI-1640 with 25 mM HEPES, 11.50 mg/L hypoxanthine, 24 mM sodium bicarbonate and 4.31 mg/mL AlbuMAX II, Invitrogen) at 37°C under 1% O₂, 5% CO₂ and 94% N₂. Complete medium was prepared using isoleucine-free RPMI (US Biological, R9014) supplemented with isoleucine (Gibco) to the standard RPMI concentration (0.38 mM). Before the assay, parasites were synchronized to ring stage with 5% sorbitol (twice, in subsequent cycles), washed four times in isoleucine-free RPMI, and diluted to ≈0.5% parasitemia using the corresponding donor RBCs in either isoleucine-free RPMI or RPMI reconstituted with isoleucine to the desired level. Growth was assessed by flow cytometry at 0, 24, 48, 72 and 96 h. Blood smears were collected at the same timepoints and stained with Hemacolor. For parasitemia, samples were washed once in PBS/0.5% BSA and resuspended in PBS/0.5% BSA with Vybrant DyeCycle Green (1:1000, Invitrogen V35004), incubated 15 min at room temperature, and acquired on a Cytek Northern Lights.

For HbF quantification, samples were collected at assay setup, fixed overnight at 4°C in 4% PFA/0.0075% glutaraldehyde in PBS, permeabilized with 0.1% Triton X-100 (20 min), stained with anti-HbF–PE (Invitrogen MHFH04; 1:100 in PBS/0.5% BSA) 30 min at 4°C, washed six times, incubated with Vybrant DyeCycle Violet (Invitrogen V35003; 1:5000) 15 min at room temperature, and acquired on a Cytek Northern Lights. Data were analyzed in FlowJo v10.

#### Parasitemia and fold-change quantification

Flow cytometry data were analyzed to determine parasitemia as the percentage of infected erythrocytes (Vybrant DyeCycle Green-positive) among total gated single cells at each timepoint. For each sample, growth was expressed as the fold change in parasitemia relative to its own t0 value, matched on blood donor, technical replicate and isoleucine concentration. The two technical replicates per condition were averaged to a single value per biological replicate. Uninfected control wells were used to set gating thresholds and were excluded from growth analyses. Data were processed in Python using pandas and NumPy.

#### Developmental stage scoring

Parasite developmental stages were scored from Hemacolor-stained blood smears and classified as ring, young trophozoite, old trophozoite or schizont, counting at least 100 parasites per smear. For each smear, stage counts were expressed as the percentage of each stage among total counted parasites. Stage composition was analyzed for adult and cord blood at 0 and 381 µM isoleucine across all timepoints (three donors per blood type). In addition to relative stage composition (percentage of parasites), absolute stage abundance was calculated by multiplying each stage’s fractional composition by the total parasitemia fold change, so that the stacked composition reflects the size of the parasite population from which it was drawn.

#### Statistical analysis

Analyses were performed on log₂-transformed parasitemia fold-change values. Technical and biological replicates were averaged to a single value per donor per condition before analysis (n = 3 donors per blood category). All tests contrast 0 µM with the physiological 70 µM isoleucine condition. Within each blood category, endpoint (96 h) log_2_ fold change was compared between the two isoleucine conditions by paired two-tailed t-test. To test whether the effect of isoleucine differed between adult and fetal blood while accounting for the repeated-measures structure, a linear mixed-effects model was fitted by maximum likelihood to log₂ fold change across the full time-course, with fixed effects for blood type, isoleucine concentration, time and their interactions and a random intercept for donor. Time was entered as a continuous variable scaled to the 96-h endpoint, so the blood type x isoleucine x time interaction tested whether isoleucine altered the rate of parasite growth differently in adult and cord blood. The corresponding two-way term estimates the contrast at t = 0, where cultures were seeded at matched parasitemia by design. Analyses were performed in Python using SciPy (paired t-tests) and statsmodels (mixed-effects model).

### Statistical analysis and software

All analyses and figures were produced in Python 3.12. Phylogenetic generalized least squares, maximum-likelihood estimation of Pagel’s λ, the phylogenetic simulation test and the model comparisons were implemented directly using SciPy and DendroPy rather than a dedicated comparative-methods package. Figures were drawn in Matplotlib. Package versions are given in the environment files in the Extended Data^14^.

## DATA AND CODE AVAILABILITY

Processed data tables underlying all figures, together with the curated species, host-preference and gene-assignment tables and the full lists of RefSeq accessions for the genomes analyzed in this study, are available in the Extended Data^14^. The RNA sequencing datasets are available at NCBI under BioProjects PRJNA1502948^10^, PRJNA796320^49^, PRJNA752401^50^.

All custom code used in this paper is archived in the Extended Data^14^. Claude (Anthropic) and Gemini (Google) were used to assist with code, plotting, and data-consistency checks, all of which were reviewed and verified by the authors.

## REFERENCES

1. Mazel, D. & Marlière, P. Adaptive eradication of methionine and cysteine from cyanobacterial light-harvesting proteins. Nature 341, 245–248 (1989).

2. Shenhav, L. & Zeevi, D. Resource conservation manifests in the genetic code. Science 370, 683–687 (2020).

3. Soghigian, J. et al. Phylogenomics reveals the history of host use in mosquitoes. Nat Commun 14, 6252 (2023).

4. Clements, A. N. Vitellogenesis. in The Biology of Mosquitoes, Volume 1: Development, Nutrition and Reproduction 360–379 (1992). doi:10.1079/9780851993744.0020.

5. Merritt, R. W., Dadd, R. H. & Walker, E. D. Feeding Behavior, Natural Food, and Nutritional Relationships of Larval Mosquitoes. Annual Review of Entomology 37, 349–374 (1992).

6. Briegel, H. Metabolic relationship between female body size, reserves, and fecundity of *Aedes aegypti*. Journal of Insect Physiology 36, 165–172 (1990).

7. Stein, W. H., Kunkel, H. G., Cole, R. D., Spackman, D. H. & Moore, S. Observations on the amino acid composition of human hemoglobins. Biochimica et Biophysica Acta 24, 640–642 (1957).

8. Harrison, R. E., Brown, M. R. & Strand, M. R. Whole blood and blood components from vertebrates differentially affect egg formation in three species of anautogenous mosquitoes. Parasites Vectors 14, 119 (2021).

9. Chang, Y.-Y. H. & Judson, C. L. The role of isoleucine in differential egg production by the mosquito *Aedes aegypti* Linnaeus (Diptera: Culicidae) following feeding on human or guinea pig blood. Comparative Biochemistry and Physiology Part A: Physiology 57, 23–28 (1977).

10. Houri-Zeevi, L., et al. Convergent evolutionary loss of chemosensory and blood-feeding pathways in non-blood-feeding mosquitoes. *bioRxiv*, DOI pending (2026a).

11. Liu, J., Istvan, E. S., Gluzman, I. Y., Gross, J. & Goldberg, D. E. Plasmodium falciparum ensures its amino acid supply with multiple acquisition pathways and redundant proteolytic enzyme systems. Proceedings of the National Academy of Sciences 103, 8840–8845 (2006).

12. Bryk, A. H. & Wiśniewski, J. R. Quantitative Analysis of Human Red Blood Cell Proteome. J. Proteome Res. 16, 2752–2761 (2017).

13. Anderson, N. L. & Anderson, N. G. The Human Plasma Proteome: History, Character, and Diagnostic Prospects*. Molecular & Cellular Proteomics 1, 845–867 (2002).

14. Houri-Zeevi, L., Kampmann, M., Duraisingh, M. T. & Vosshall, L. B. Extended data for ‘Isoleucine absence from adult human hemoglobin is mirrored in mosquito proteomes and limits malaria parasite growth’. 10.5281/zenodo.22699764.

15. Sueoka, N. Directional mutation pressure and neutral molecular evolution. Proceedings of the National Academy of Sciences 85, 2653–2657 (1988).

16. Duret, L. & Galtier, N. Biased gene conversion and the evolution of mammalian genomic landscapes. Annu Rev Genomics Hum Genet 10, 285–311 (2009).

17. Akashi, H. & Gojobori, T. Metabolic efficiency and amino acid composition in the proteomes of Escherichia coli and Bacillus subtilis. Proceedings of the National Academy of Sciences 99, 3695–3700 (2002).

18. Clements, A. N. Adult digestion. in The Biology of Mosquitoes, Volume 1: Development, Nutrition and Reproduction 272–291 (1992). doi:10.1079/9780851993744.0014.

19. Sankaran, V. G. & Orkin, S. H. The Switch from Fetal to Adult Hemoglobin. Cold Spring Harb Perspect Med 3, a011643 (2013).

20. Fanali, G. et al. Human serum albumin: From bench to bedside. Molecular Aspects of Medicine 33, 209–290 (2012).

21. Vidarsson, G., Dekkers, G. & Rispens, T. IgG Subclasses and Allotypes: From Structure to Effector Functions. Front. Immunol. 5, (2014).

22. Francis, S. E., Sullivan, D. J., Goldberg & E, D. Hemoglobin Metabolism in the Malaria Parasite *Plasmodium Falciparum*. Annual Review of Microbiology 51, 97–123 (1997).

23. Martin, R. E. & Kirk, K. Transport of the essential nutrient isoleucine in human erythrocytes infected with the malaria parasite *Plasmodium falciparum*. Blood 109, 2217–2224 (2006).

24. Babbitt, S. E., et al. *Plasmodium falciparum* responds to amino acid starvation by entering into a hibernatory state. Proceedings of the National Academy of Sciences 109, E3278–E3287 (2012).

25. Bard, H. The postnatal decline of hemoglobin F synthesis in normal full-term infants. J Clin Invest 55, 395–398 (1975).

26. Fried, M. & Duffy, P. E. Adherence of *Plasmodium falciparum* to Chondroitin Sulfate A in the Human Placenta. Science 272, 1502–1504 (1996).

27. Rogerson, S. J., Hviid, L., Duffy, P. E., Leke, R. F. G. & Taylor, D. W. Malaria in pregnancy: pathogenesis and immunity. Lancet Infect Dis 7, 105–117 (2007).

28. Trager, W. & Jensen, J. B. Human Malaria Parasites in Continuous Culture. Science 193, 673–675 (1976).

29. Pasvol, G., Weatherall, D. J. & Wilson, R. J. M. Effects of fetal haemoglobin on susceptibility of red cells to Plasmodium falciparum. Nature 270, 171–173 (1977).

30. Archer, N. M., Petersen, N. & Duraisingh, M. T. Fetal hemoglobin does not inhibit *Plasmodium falciparum* growth. Blood Adv 3, 2149–2152 (2019).

31. Archer, N. M., Pasvol, G., Wilson, I. & Duraisingh, M. T. Does Fetal Hemoglobin inhibit the malarial parasite *Plasmodium falciparum*? Am J Hematol 97, E325–E327 (2022).

32. Gavulic, A. E. et al. Fetal hemoglobin levels in premature newborns. Should we reconsider transfusion of adult donor blood? Journal of Pediatric Surgery 56, 1944–1948 (2021).

33. McLean, K. J. & Jacobs-Lorena, M. The response of *Plasmodium falciparum* to isoleucine withdrawal is dependent on the stage of progression through the intraerythrocytic cell cycle. Malar J 19, 147 (2020).

34. Marreiros, I. M. et al. A non-canonical sensing pathway mediates *Plasmodium adaptation* to amino acid deficiency. Commun Biol 6, 205 (2023).

35. Elf, J., Nilsson, D., Tenson, T. & Ehrenberg, M. Selective Charging of tRNA Isoacceptors Explains Patterns of Codon Usage. Science 300, 1718–1722 (2003).

36. Liu, L. et al. Toward life with a 19-amino acid alphabet through generative artificial intelligence design. Science 392, eaeb5171 (2026).

37. Kwiatkowski, D. P. How Malaria Has Affected the Human Genome and What Human Genetics Can Teach Us about Malaria. The American Journal of Human Genetics 77, 171–192 (2005).

38. Winegard, T. C. The Mosquito: A Human History of Our Deadliest Predator. (Dutton, New York, 2019).

39. Allison, A. C. Protection Afforded by Sickle-cell Trait Against Subtertian Malarial Infection. Br Med J 1, 290–294 (1954).

40. Frangoul, H. et al. CRISPR-Cas9 Gene Editing for Sickle Cell Disease and β-Thalassemia. New England Journal of Medicine 384, 252–260 (2021).

41. Frangoul, H. et al. Exagamglogene Autotemcel for Severe Sickle Cell Disease. New England Journal of Medicine 390, 1649–1662 (2024).

42. Piel, F. B. et al. Global epidemiology of sickle haemoglobin in neonates: a contemporary geostatistical model-based map and population estimates. The Lancet 381, 142–151 (2013).

43. Weinberg, E. D. Nutritional Immunity: Host’s Attempt to Withhold Iron From Microbial Invaders. JAMA 231, 39–41 (1975).

44. Hood, M. I. & Skaar, E. P. Nutritional immunity: transition metals at the pathogen–host interface. Nat Rev Microbiol 10, 525–537 (2012).

45. Kok, G. et al. Isoleucine-to-valine substitutions support cellular physiology during isoleucine deprivation. Nucleic Acids Res 53, gkae1184 (2025).

46. Ebel, E. R., Kuypers, F. A., Lin, C., Petrov, D. A. & Egan, E. S. Common host variation drives malaria parasite fitness in healthy human red cells. eLife 10, e69808 (2021).

47. Emms, D. M., Liu, Y., Belcher, L., Holmes, J. & Kelly, S. OrthoFinder: improved phylogenetic orthology inference with enhanced accuracy and scalability. Nat Methods 23, 1327–1333 (2026).

48. Garland, T., Dickerman, A. W., Janis, C. M. & Jones, J. A. Phylogenetic Analysis of Covariance by Computer Simulation. Systematic Biology 42, 265–292 (1993).

49. Venkataraman, K. et al. Two novel, tightly linked, and rapidly evolving genes underlie *Aedes aegypti* mosquito reproductive resilience during drought. eLife 12, e80489 (2023).

50. Martinson, E. O., Chen, K., Valzania, L., Brown, M. R. & Strand, M. R. Insulin-like peptide 3 stimulates hemocytes to proliferate in anautogenous and facultatively autogenous mosquitoes. J Exp Biol 225, jeb243460 (2022).

51. Martin, M. Cutadapt removes adapter sequences from high-throughput sequencing reads. EMBnet.journal 17, 10–12 (2011).

52. Bray, N. L., Pimentel, H., Melsted, P. & Pachter, L. Near-optimal probabilistic RNA-seq quantification. Nat Biotechnol 34, 525–527 (2016).

53. Soneson, C., Love, M. I. & Robinson, M. D. Differential analyses for RNA-seq: transcript-level estimates improve gene-level inferences. F1000Res 4, 1521 (2015).

54. Babraham Bioinformatics -FastQC A Quality Control tool for High Throughput Sequence Data. https://www.bioinformatics.babraham.ac.uk/projects/fastqc/.

55. Goldman, O. V. et al. Supplementary data for ‘A single-nucleus transcriptomic atlas of the adult *Aedes aegypti mosquito*’ (Goldman et al., 2025). https://doi.org/10.5281/zenodo.17382236 (2025) doi:10.5281/zenodo.17382236.

56. Katoh, K. & Standley, D. M. MAFFT multiple sequence alignment software version 7: improvements in performance and usability. Mol Biol Evol 30, 772–780 (2013).

